# PrP turnover in vivo and the time to effect of prion disease therapeutics

**DOI:** 10.1101/2024.11.12.623215

**Authors:** Taylor L Corridon, Jill O’Moore, Alyssa Seerley, Daniel A. Sprague, Yuan Lian, Vanessa Laversenne, Briana Noble, Nikita G Kamath, Fiona E Serack, Abdul Basit Shaikh, Brian Erickson, Craig Braun, Kendrick DeSouza-Lenz, Michael Howard, Nathan Chan, Walker S Jackson, Andrew G Reidenbach, Deborah E Cabin, Sonia M Vallabh, Andrea Grindeland Panter, Nina Oberbeck, Hien T Zhao, Eric Vallabh Minikel

## Abstract

PrP lowering is effective against prion disease in animal models and is being tested clinically. Therapies in the current pipeline lower PrP production, leaving pre-existing PrP to be cleared according to its own half-life. We hypothesized that PrP’s half-life may be a rate-limiting factor for the time to effect of PrP-lowering drugs, and one reason why late treatment of prion-infected mice is not as effective as early treatment. Using isotopically labeled diet with targeted mass spectrometry, as well as antisense oligonucleotide treatment followed by timed PrP measurement, we estimate a half-life of 5-6 days for PrP in the brain. PrP turnover is not affected by over-or under-expression. Mouse PrP and human PrP have similar turnover rates measured in wild-type or humanized knock-in mice. CSF PrP appears to mirror brain PrP in real time in rats. PrP in the colon is readily quantifiable and has a half-life just slightly shorter than in brain. An under-expressed pathogenic mutant PrP, corresponding to D178N in humans, exhibits an accelerated turnover rate. Our data may inform the design of both preclinical and clinical studies of PrP-lowering drugs.

**Author Summary:** Prion disease is a fatal brain disease caused by misfolding of the prion protein (PrP). Emerging therapies for prion disease seek to reduce the amount of PrP produced in the brain in order to delay onset of disease or slow progression. Mouse studies have shown that if these therapies are initiated too late, their benefit is limited or they may not help at all. Here we measure the half-life of PrP in the mouse brain, and find that it is about 5 days. When drugs are used to lower PrP by cutting up the RNA that encodes PrP, the RNA drops rapidly while the protein lags behind, and does not reach its minimum level until 4 weeks after the drug is dosed. This half-life is about the same regardless of the species of PrP (mouse or human) and whether or not the brain is infected with prions. Cerebrospinal fluid appears to reflect the real-time levels of brain PrP with no appreciable lag. PrP can be measured in colon, which may be useful in animal studies of systemic drugs to lower PrP. PrP turns over more quickly in the presence of a pathogenic genetic variant, the equivalent of the human D178N variant. These findings suggest that clinical trials can monitor PrP in cerebrospinal fluid to look at drug activity, but should plan timepoints far enough post-dose to account for PrP’s rate of turnover, and should focus on patients who will survive long enough to benefit from the drug.

## Introduction

Pharmacologic lowering of prion protein (PrP) delays onset and slows progression of prion disease in animal models [1–3], consistent with PrP as the substrate for prion misfolding and the pivotal molecule in progression of this rapid neurodegenerative disease [4,5]. Inspired by this finding, a PrP-lowering antisense oligonucleotide (ASO) is now in a Phase I clinical trial (NCT06153966), with additional PrP-lowering modalities in preclinical development [6].

Prion disease typically presents as a rapidly progressive dementia [7]. With the median patient in prior clinical trials surviving just ∼2 months from randomization [8,9], the time to effect of prion disease therapeutics could be a critical determinant of efficacy in the symptomatic population. In mouse models, PrP lowering is most effective when treatment is administered early in the silent incubation period (<78 days post-inoculation or dpi) [1]. Treatment after frank symptoms emerge (132 – 143 dpi) has extended survival primarily by increasing the time that animals are sick, without reverting any symptoms already accumulated [1,2], and only a subset of late-treated animals benefit, while others succumb to disease on a similar timeframe as untreated animals [1]. One explanation is simply that PrP lowering cannot reverse existing neuronal loss. However, the observation that efficacy is limited even very late pre-symptomatic timepoints (105 – 120 dpi) [1] led us to speculate that PrP turnover may be another factor limiting the efficacy of late treatment. ASOs target the PrP RNA for cleavage and degradation by RNase H1 [1,10,11], suppressing new PrP synthesis but leaving pre-existing PrP to be degraded according to its own half-life. PrP turns over rapidly in cultured cells [12,13], but in vivo, reports are conflicting. A study using oral doxycycline to suppress expression of PrP under a Tet-off transgene determined the half-life of normally folded cellular PrP (PrP^C^) to be just 0.75 days in the brain [14], while 2 mass spectrometry studies of mice fed isotopically labeled chow determined half-life estimates of 4.95 or 5.02 days in the brain [15,16]. Given that prion disease has heterogeneous subtypes with distinct rates of progression [7], the difference between a half-life of <1 day versus 5 days would have a dramatic impact on the inclusion criteria needed to select for patients likely to have time to benefit from a PrP-lowering drug in clinical trials.

Here we set out to determine the half-life of PrP in brain, as well as to answer several related questions. Because some PrP-lowering drugs in development are expected to have systemic activity [6], we sought to identify an organ or tissue that could be used as a proxy for peripheral target engagement in preclinical models, and further to determine the PrP half-life and therefore timeline on which target engagement can be observed in such a tissue. To mitigate translational risk due to amino acid sequence differences, depth of target suppression, or disease state, we sought to determine whether PrP half-life differs between human versus mouse PrP, in heterozygous knockout versus overexpressing animals, or in prion-infected versus naïve animals. Because cerebrospinal fluid (CSF) is being used as a sampling compartment to reflect on brain PrP [5,17], we also sought to determine the timeframe on which brain target engagement can be read out in CSF. Finally, because pathogenic variants in neurodegenerative disease-associated proteins can impact protein half-life [18], we sought to determine whether the pathogenic D178N variant, which results in reduced PrP expression [19,20], alters PrP’s half-life.

## Results

We sought to identify a peripheral tissue in which we could quantify PrP. Analysis of human *PRNP* RNA expression data from Genotype-Tissue Expression project (GTEx v8) [21] revealed that after brain and sciatic nerve, colon was the next tissue with the highest *PRNP* expression (Figure 1A). Similarly in mouse, colon ranked third after brain and esophagus for Prnp RNA expression [22] (Figure 1B). We dissected 15 organs from 1 wild-type and 1 PrP knockout mouse and analyzed them by Western blot. Brain exhibited far more PrP than any peripheral organ examined, but a strong band was identified centered at the expected molecular weight (∼37 kDa) in colon, with weaker bands in stomach, quadriceps, heart, femur, spleen, and uterus, and faintly detectable bands in lung, lymph node, and skin. PrP was not detectable in liver, kidney, whole blood, or plasma (Figure 1C). Although homogenization efficiency and total protein loading varied between organs, Coomassie analysis revealed that the WT and KO animals were similarly loaded for any given organ (Figure 1C). When the same tissues were analyzed at a 1:100 wt/vol final dilution by our in-house ELISA^17^ with the EP1802Y/8H4 antibody pair, all tissues besides brain were near the lower limit of quantification (LLQ), with many reading above LLQ in the knockout animals, presumably due to matrix effects (Figure 1D). Of any organ where the knockout tissue read out at LLQ, colon exhibited the strongest PrP signal in the wild-type animal (Figure 1D). Colon and 4 tissues with weaker signal were re-analyzed at a 1:25 wt/vol final dilution, yielding higher signal and confirming colon as the strongest tissue at 5-fold above LLQ (Figure 1E). Further assay development identified the best conditions for ELISA detection of colon PrP and established stability parameters for colon samples in this assay (Figure S1). These results led us to nominate colon as a proxy tissue for monitoring PrP turnover and knockdown in animal experiments (see Discussion).

**Figure 1.**
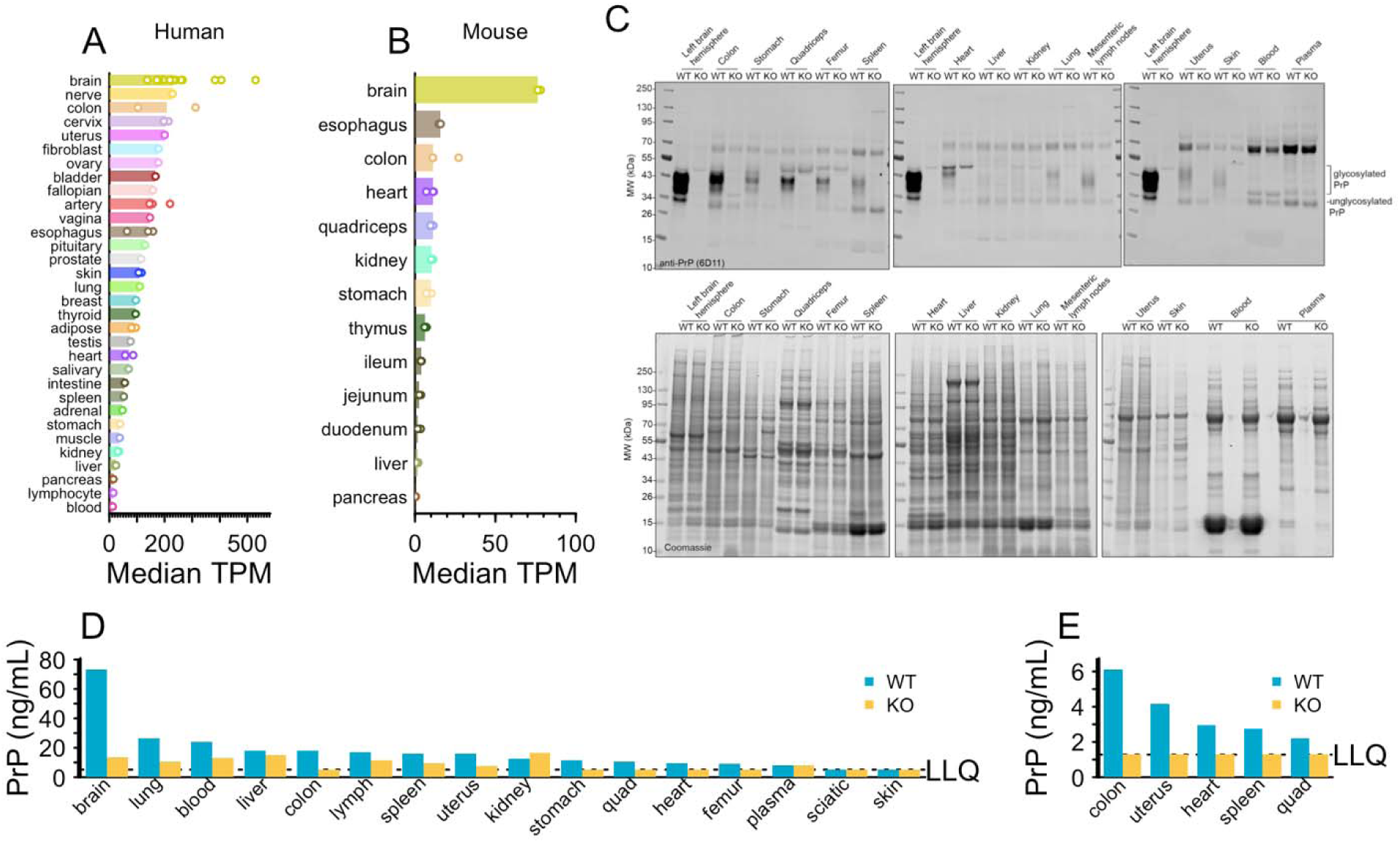
Nomination of colon as a tissue for peripheral PrP quantification. **A)** PRNP RNA expression in transcripts per million (TPM) in human tissues according to GTEx v8 public data. Each sub-tissue (e.g. brain – cerebellum) is represented by one point as the median TPM across all samples for that tissue, and each tissue (e.g. brain) is represented by one bar as the median of those medians. **B)** Prnp RNA expression in TPM in mouse tissues according to Söllner et al [22]. Each animal is represented by one point for individually measured TPM, and each tissue is represented by one bar as the median of the animals. **C)** Western blot (top) and Coomassie (bottom) of organs all from the same 1 WT and 1 KO animal, 6D11 anti-PrP antibody, see Methods for details. **D)** Organs tested by PrP ELISA at a 1:100 final dilution (10% homogenates at a further 1:10). PrP ELISA as reported except using double the detection mAb concentration (0.5 µg/mL instead of 0.25 µg/mL). **E)** Organs tested by PrP ELISA at a 1:25 final dilution (10% homogenates at a further 1:2.5). PrP ELISA as reported except using double the detection mAb concentration (0.5 µg/mL instead of 0.25 µg/mL). See Figure S1 for further assay development.

We next sought to use targeted MS of PrP tryptic peptides (Figure 2A) to measure turnover in both brain and colon. Initially we focused solely on the peptide VVEQMCVTQYQK (mouse PrP residues 208-219; hereafter abbreviated VVEQ), the most readily quantified of any PrP tryptic peptide [23], in both brain and colon (Figure S2). We fed wild-type mice with ^13^C_6_ lysine chow, sacrificed them at 0, 2, 4, 6, or 8 days — a range of timepoints around the hypothesized half-life of PrP based on prior mass spectrometry studies [15,16] — measured VVEQ by targeted mass spectrometry, and quantified the percent labeled as the ratio of heavy peptide to heavy plus light. Due to lower overall PrP abundance (Figure S2), the LLQ in colon occurred at 13.3% labeling versus 2.5% for brain; nonetheless, heavy peptide was above LLQ in most samples by day 2, and in all samples on days 4-8 (Figure 2B). Heavy labeled peptide accumulated much more quickly in colon than in brain, with the two tissues reaching 49.3% and 20.5% respectively by day 8 (Figure 2B), potentially suggesting a shorter half-life in colon than in brain.

**Figure 2.**
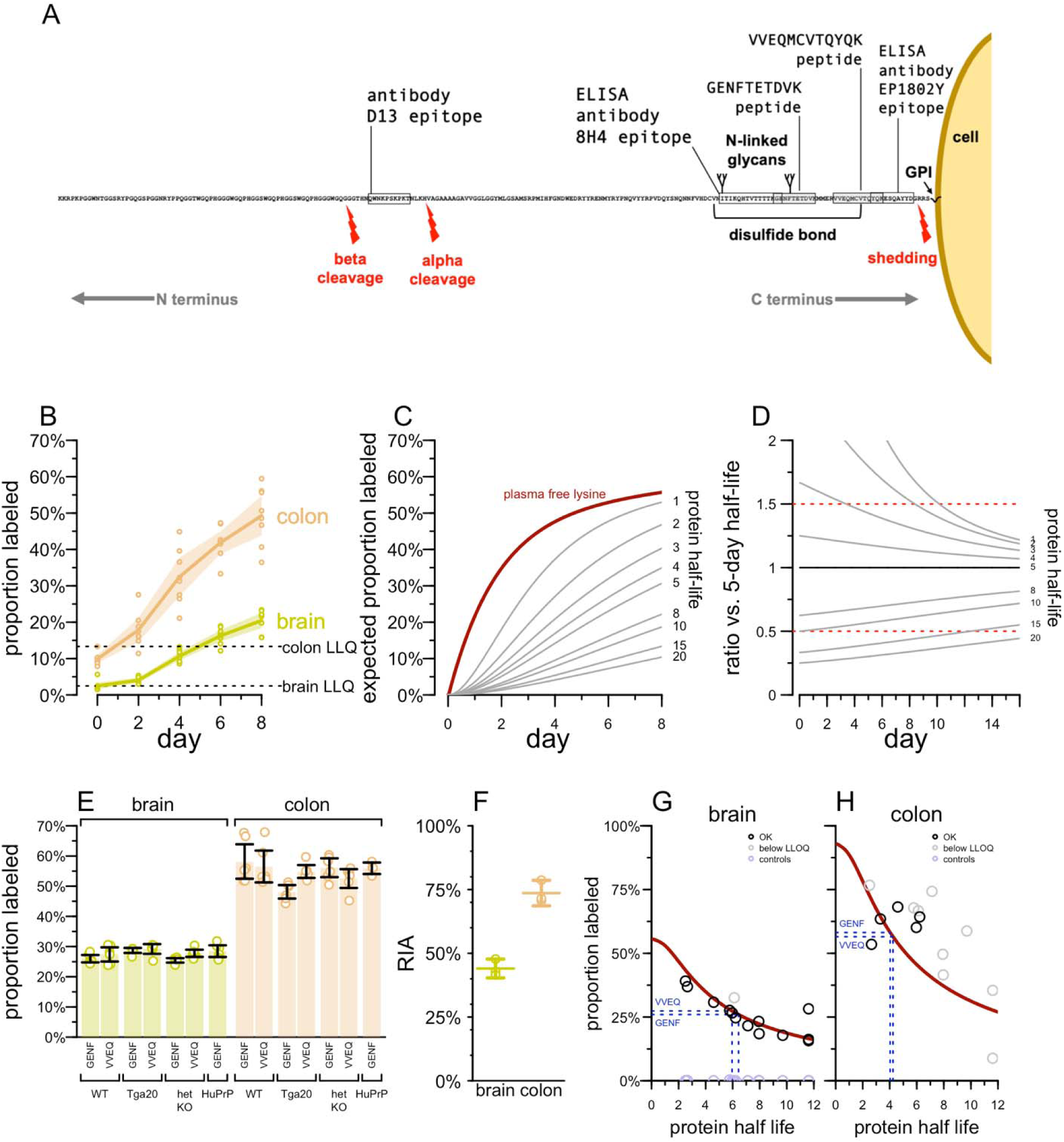
Determination of PrP half-life by targeted mass spectrometry. **A)** Diagram of mature, post-translationally modified mouse PrP. Adapted from a CC BY licensed diagram we have previously released (https://github.com/ericminikel/prp_mrm/), see Minikel & Kuhn et al for details [23]. Indicated are the positions of peptides measured here in mass spectrometry, the epitopes of antibodies used in our in-house ELISA as well as D13, the antibody used for Western blot PrP quantification and conformation-dependent immunoassay by Safar et al [14]. **B)** Accumulation of ^13^C_6_ label from chow in the VVEQ peptide in mouse brain and colon. **C)** The best fit to the proportion of plasma free lysine empirically found to be ^13^C_6_ labeled, as reported by Fornasiero (maroon), and the proportion of peptide expected to be labeled over time as a function of half-life (shown in days on the right side). See Methods > Labeled peptide accumulation models for details. **D)** The ratio of proportion labeled (C) for a peptide of each half-life compared to a peptide of 5-day half-life. Ratios of 0.5 and 1.5 are arbitrary landmarks highlighted to orient the eyes to a straight horizontal line. **E)** The proportion of PrP peptides VVEQ and GENF that are ^13^C_6_ labeled after 8 days as a function of mouse genotype. All differences are non-significant at Bonferroni-corrected P > 0.05, 2-sided T-test. **F)** Relative isotope abundance (RIA), in other words, the mean ratio of heavy to light lysine inferred to have been available over the 8-day labeling period, by tissue. Each point is one animal (N=3 wild-type C57BL/6N per tissue). Segments represent means and error bars 95% confidence intervals per tissue. **G)** Proportion labeled in brain (y axis) versus brain half-life previously reported by Fornasiero for all measured non-PrP peptides (circles); black = measured peptide signal above LLQ, gray = below LLQ. The maroon line represents the expected proportion labeled after 8 days as a function of half-life, based on the plasma free lysine model from (C). The horizontal blue lines represent the proportion labeled observed for the two PrP peptides, and their vertical projection from the maroon curve down to the x axis represents the estimation of half-life from those proportion labeled measurements. **G)** As in (F), but for colon. The maroon line uses the plasma free lysine model from (C) adjusted based on the ratio of empirical RIA in colon compared to brain from (F).

In order to interpret these data, we considered implications of the mathematical model for heavy label accumulation presented by Fornasiero et al [16]. Fornasiero measured the proportion of free lysine in mouse plasma that was labeled, and fit a model represented by the maroon line (Figure 2C). In this model, the proportion of lysine that is heavy labeled rises rapidly initially as dietary lysine becomes bioavailable, but then increases more slowly, reaching 55.7% by day 8, because the labeled dietary lysine is in competition with unlabeled lysine made available by catabolism of endogenous proteins. Because only a portion of free lysine is labeled, calculating the expected proportion of a peptide labeled as a function of its protein’s half-life revealed that according to this model, peptides from a protein with a 5-day half-life would be just 30.0% labeled by day 8, even though 62.1% of the protein would turn over in this time (Figure 2D).

With this in mind, we considered how many days of labeled chow consumption would best discriminate between shorter and longer half-lives. This analysis revealed a tradeoff: the theoretical difference between proportion labeled for a quick turnover protein and a slow turnover protein is maximized at early timepoints when the overall proportion labeled is still low enough that the precision of measurement near LLQ could be limiting. At later timepoints, the proportion labeled is higher, mitigating LLQ concerns, but the theoretical proportion labeled is less different. For discriminating half-lives near 5 days, an 8-day labeled chow experiment appeared to present a reasonable compromise between these tradeoffs.

To replicate and extend our results, we performed a multiplex targeted MS assay using VVEQ, another PrP peptide GENFTETDVK (mouse PrP residues 194-203; hereafter abbreviated GENF), and a sampling of peptides from proteins whose brain half-lives as determined by Fornasiero [16] ranged from 2.5 to 11.6 days, to serve as controls. Serial dilution of ^13^C_6_ ^15^N_2_ lysine synthetic peptides for this assay identified lower limits of quantification (LLQ) for each peptide; the mean heavy peptide area found in wild-type mice after 8 days of labeled chow was above LLQ for 17 peptides in brain and for 8 in colon, indicating the suitability of these peptides for this purpose (Figure S3).

We utilized multiple mouse lines (Table 1; Figure S4-S5) to determine the impact of PrP amino acid sequence and expression level on half-life. For the multiplex MS assay, we fed wild-type, heterozygous PrP knockout, transgenic humanized (Tg25109; human PrP 129M), and transgenic overexpressing (Tga20 mouse PrP) mice with ^13^C_6_ lysine chow, sacrificed them at 8 days, and analyzed their brains by mass spectrometry. For either PrP peptide, measured in either tissue, the proportion labeled was not significantly different from wild-type for any genotype (Figure 2E).

**Table 1.** Mouse lines used in this study. All PrP expression levels are measured in whole hemisphere, either by our in-house ELISA [17] or by the IQ Proteomics MRM assay described herein, and are shown normalized to wild-type animals (1.0x = wild-type epxression). *Maintained on a background of homozygous ZH3/ZH3 PrP knockout. **MoPrP-A refers to the mouse reference genome PrP sequence as found in C57BL/6N and most other commonly used mouse strains (as opposed to the MoPrP-B allele, containing the two substitutions L108F and V189T, found in certain strains [24]). ***The 3F4 epitope is two amino acid changes from the MoPrP-A sequence: L108M + V111M. ****Mean value obtained for 2 peptides in young and aged ki-3F4-FFI mice in Figure 4 of this study.

| Name | PrP amino acid sequence | Expression level (fold wild-type) | PrP quantification method | Genomic location | Reference |
| --- | --- | --- | --- | --- | --- |
| ZH3/+ | MoPrP-A** | 0.5x | <u>ELISA</u> | *Prnp* | [25] |
| Tg25109* | HuPrP 129M | 1.1x | <u>ELISA</u> | *Frdm6/Tmx1* | [26] |
| Ki817 | HuPrP 129V | 1.0x | <u>ELISA</u> | *Prnp* | This study<br>Fig S4 |
| Tga20* | MoPrP-A | 2.4x | <u>ELISA</u> | *Ptcra* | [27], this<br>study Fig<br>S5 |
| ki-3F4-WT | MoPrP-3F4*** | 1.0x | MRM | *Prnp* | [28] |
| ki-3F4-FFI | MoPrP-3F4-D177N | 0.4x**** | MRM | *Prnp* | [28], this<br>study Fig<br>4 |

Across PrP peptides and all mouse genotypes, the proportion labeled in colon was approximately double that in brain (54.6% vs. 27.6%, Figure 2E). Indeed, even the plasma free lysine concentration is not expected to 54.6% labeled until day 7 (Figure 2C), which would mean that, instantaneous under the plasma free lysine model, PrP turnover in colon would need to be virtually in order to account for this high a percent labeled. We considered instead the possibility that the colon absorbs lysine directly from the diet, bypassing the bloodstream. In order to empirically determine the availability of free lysine in each tissue, we employed limited trypsin digestion followed by data-independent acquisition (DIA) proteomics on matched pairs of brain and colon samples from 3 wild-type C57BL/6N mice fed labeled chow for 8 days (Figure 2F, S6A-B). This allowed us to quantify N=833 double-lysine peptides owing to missed trypsin cleavages, from N=596 distinct proteins. Regardless of protein half-life, double-lysine peptides with at least one heavy lysine are necessarily nascent, synthesized during the 8 days of labeled diet, while the ratio of peptides with one light and one heavy lysine (LH) to those with both heavy lysines (HH) provides information on the mean relative isotope abundance (RIA), or the proportion of free lysine that was heavy at the time these various proteins were synthesized [29,30] (see Methods). The point estimate of RIA obtained by this method varied among peptides across the proteome, yet the distribution was highly consistent within each tissue between different animals (Figure S6A) and was also consistent within each tissue between different bins of protein abundance (Figure S6B). Overall, the estimated RIA was 44.0% in brain (Figure 2F), in good agreement with the 41.8% mean expected over the 8-day period under the plasma free lysine model (Figure 2C). RIA was 73.6% in colon (Figure 2F), 1.67 times higher than brain, confirming our suspicion that colon had more ready access to dietary labeled lysine than brain did.

When we plotted, for each peptide from various proteins, the proportion labeled in wild-type mouse brain at day 8 versus the half-life reported by Fornasiero (Figure 2G), we found excellent agreement with the theoretical proportion labeled expected based on the plasma free lysine curve (Figure 2D). Projecting the proportion labeled in brain in WT mice for the two PrP peptides (26.0% and 27.3% for GENF and VVEQ respectively) onto this curve (blue dashed lines) yielded estimates of 6.4 and 6.0 days respectively. Control mice fed unlabeled chow categorically had percent labeled at <0.5%, confirming specificity of the assay. In colon (Figure 2H), we adjusted the free lysine model to account for the different RIA determined empirically in this tissue (Figure 2F). While many of the proteins with a brain half-life reported by Fornasiero were below LLQ in colon, those that were quantifiable agreed reasonably well with the theoretical proportion labeled expected based on the adjusted plasma free lysine curve.

Projecting the proportion labeled in colon in WT mice for the two PrP peptides (58.2% and 56.5% for GENF and VVEQ respectively) onto this curve (blue dashed lines) yielded estimates of 4.0 and 4.2 days respectively, just slightly lower than the half-life estimates in brain.

We also sought to determine PrP’s half-life by an orthogonal method. We dosed naïve wild-type mice with 500 µg PrP-lowering active ASO 6 [1] (Table 2) by intracerebroventricular (ICV) injection at day 0 and then performed serial sacrifice to measure *Prnp* RNA and PrP protein in whole hemispheres at various timepoints post-dose (Figure 3A). Maximal RNA suppression was achieved within 3 days, while protein lagged, reaching its nadir at 28 days (Figure 3A). When we fit an exponential decay curve to the data, we obtained a half-life estimate of 4.8 days.

**Figure 3.**
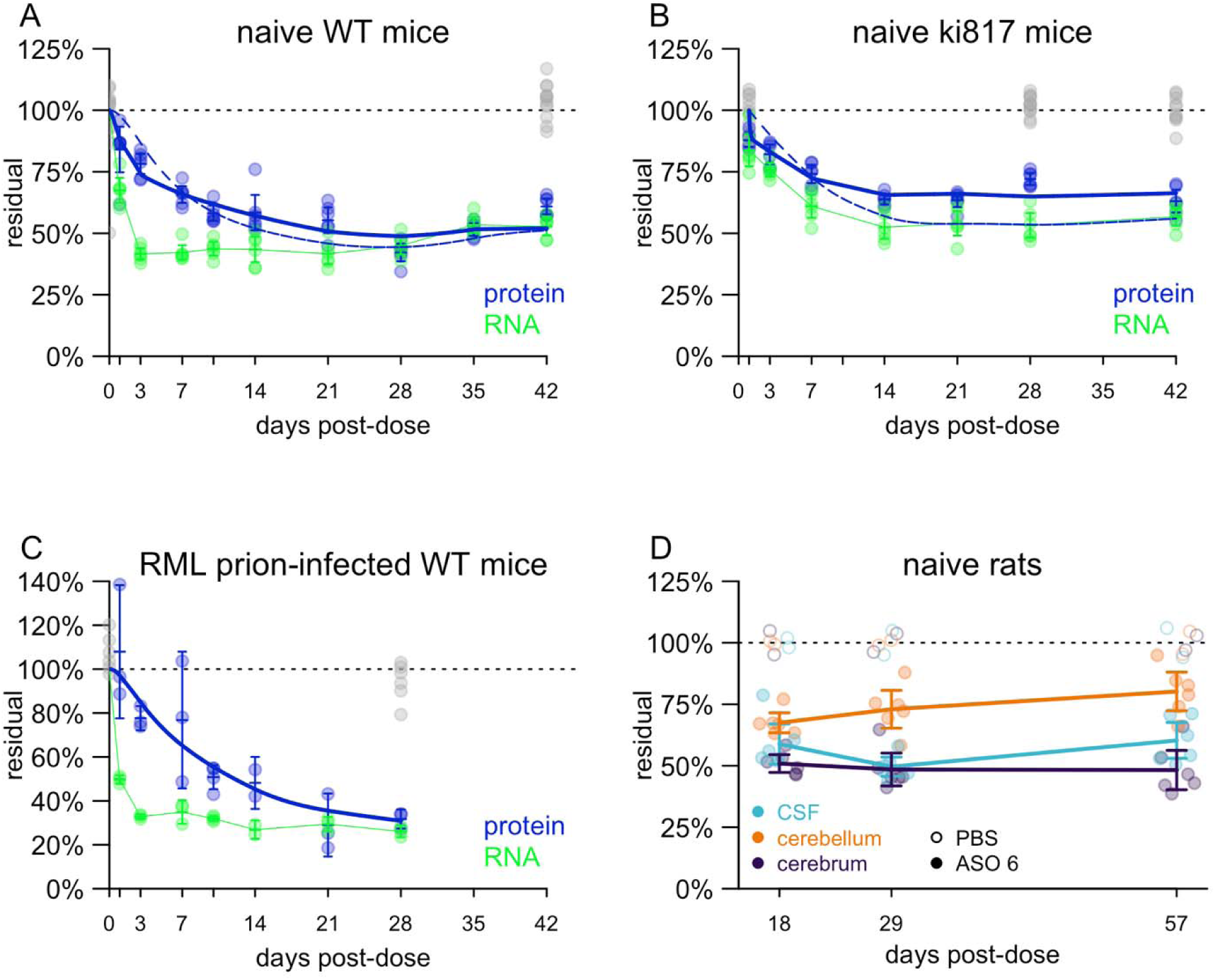
Determination of PrP half-life by ASO administration and timed sacrifice. **A)** Residual Prnp RNA and PrP protein (y axis), normalized to the mean of saline controls, at various timepoints (x axis) for wild-type mice after a single ICV dose of 500 µg active ASO 6. Each point represents a whole brain hemisphere from one animal. All measurements in saline controls (both RNA and protein) are shown in gray. For each timepoint, line segments represent means and error bars represent 95% confidence intervals. The green curve represents linearly interpolated residual RNA concentration. The dashed blue curve represents a single-rate exponential decay model fit to the data. The solid blue curve represents a mixture-of-rates fit to the data. **B)** As in (A) but for Ki817 human PrP 129V knock-in mice after a single dose 118 µg of ASO N. **C)** As in (A) but for wild-type mice infected with RML prions and treated with 300 µg ASO 6 at 105 dpi. **D)** Residual PrP in Sprague-Dawley rats treated with 1 mg of ASO 6 at day 0. Each point represents one animal, and for each timepoint, line segments represent means and error bars represent 95% confidence intervals. Long lines connect means of different timepoints.

**Table 2.** Antisense oligonucleotides used in this study. Black: unmodified DNA (2′H). Orange: 2′ methoxyethyl (2’MOE). Blue: 2′-4′ constrained ethyl (cEt). Unmarked backbone linkages: phosphorothioate (PS). Linkages marked with o = phosphodiester (PO). mC: 5-methylcytosine.

| ASO | sequence and chemistry | target | ref |
| --- | --- | --- | --- |
| ASO 6 | mCToTomCoTATTTAATGTmCAoGoTmCT | mouse/rat *Prnp* 3' UTR | [11] |
| ASO N | GTomCoAoToAoATTTTmCTTAGmCoTAmC | human/NHP *PRNP* intron | [31] |

Interestingly, while the model assumes a single rate of decay, the data do not fit such a paradigm perfectly. On one hand, we observed more knockdown at early timepoints than is explainable by a simple exponential decay model. For example, PrP protein was already down to 84% residual at day 1, when the model still predicts 98% residual. Conversely, we observed less pharmacodynamic activity at later timepoints than the model would predict: 58% residual at day 14, when the model predicts 52%. This could suggest that bulk PrP in whole brain hemisphere does not represent a single population with a uniform decay rate (see Discussion).

To confirm that there is no difference in half-life between mouse PrP and human PrP, we repeated the experiment in naïve humanized animals using a new human PrP (129V) knock-in mouse line termed Ki817 (Methods; Figure S4, S7) treated with ASO N, which is potent against *PRNP* in both human and cynomolgus macaque [31]. Targeting 50% lowering to mirror the ASO 6 experiment above, we used a dose of 118 µg, which was the median effective dose (ED_50_) estimated in mouse cortex [32]. Due to a shortage of these humanized mice, we included fewer late timepoints in this experiment, thus inadvertently biasing the model towards the early timepoints where, in our previous experiment, knockdown was deeper than predicted by exponential decay. Perhaps as a result of this bias, the estimated half-life from these data using the single-parameter model was just 2.1 days (solid blue curve, Figure 3B).

To determine whether prion infection affects the half-life of PrP, we also performed the same experiment in RML prion-infected wild-type mice with a single ICV dose of 300 µg active ASO 6 at 105 dpi (Figure 3C). This yielded a similar picture as in naïve mice, with a single-parameter half-life point estimate of 6.1 days. We note that we have not extensively tested the cross-reactivity of our PrP ELISA for PrP^Sc^; given the non-denaturing conditions of our ELISA, our assay is likely measuring primarily or exclusively PrP^C^.

To assess whether CSF PrP lags brain PrP, we used rats; the smaller CSF volume found in mice is challenging for robust PrP quantification [33]. After a single 1 mg ICV dose of active ASO 6 on day 0, rats were sacrificed at 18, 29, or 57 days post-dose and PrP was quantified in cerebrum (cortex and subcortex), cerebellum, and CSF (Figure 3D). Target engagement was deeper in cerebrum than in cerebellum at all timepoints, consistent with results from non-human primates and with the known difficulties in achieving strong ASO activity in cerebellar granule cells [31,34]. At all timepoints, the percent residual CSF PrP was in between that of cerebrum and of cerebellum, consistent with CSF reflecting some average of different brain regions. CSF PrP did not lag relative to cerebrum or cerebellum PrP, suggesting that it reflects brain PrP by 18 days post-dose, if not sooner.

Disease-causing missense variants in PrP have been shown to result in reduced PrP concentration in the CSF of asymptomatic humans at risk for prion disease [19,20] as well as in cultured cells and in the brain parenchyma of knock-in mice [28,35]. We sought to utilize a knock-in mouse model, ki-3F4-FFI, harboring the murine equivalent of the D178N variant, associated with low CSF PrP concentration in humans, to determine whether this underexpression results from accelerated turnover of PrP. Young (mean 9 weeks) or aged (mean 63 weeks) ki-3F4-FFI mice (Table 1; ref. [28]) were compared to age-matched wild-type C57BL/6N control animals as well as ki-3F4-WT, which additionally control for any impact of the 3F4 epitope as well as genetic background. All animals were fed isotopically labeled chow for 2, 4, 6, or 8 days and brains analyzed using the multiplex MS assay described above.

In young mice, the total amounts (regardless of isotopic label) of both PrP peptides (VVEQ and GENF) were reduced in the presence of the D178N substitution in ki-3F4-FFI mice, as expected (Figure 4A and S7), while decreases in abundance were not detected for any of the other proteins included in this multiplex assay. We fit empirical half-lives for every peptide in each mouse genotype; both PrP peptides exhibited significantly reduced half-lives (1.8 days for VVEQ and 2.4 days for GENF, versus 3.5 and 3.6 days respectively in control animals), while significant changes were not detected for any other protein (Figure 4B). Examination of the accumulation of heavy lysine label in these two peptides (Figure 4C-D) revealed that the proportion of peptide labeled was already above either control mouse line after just 2 days of labeled chow. The relative difference between the ki-3F4-FFI and control mice did not further increase through day 8.

**Figure 4.**
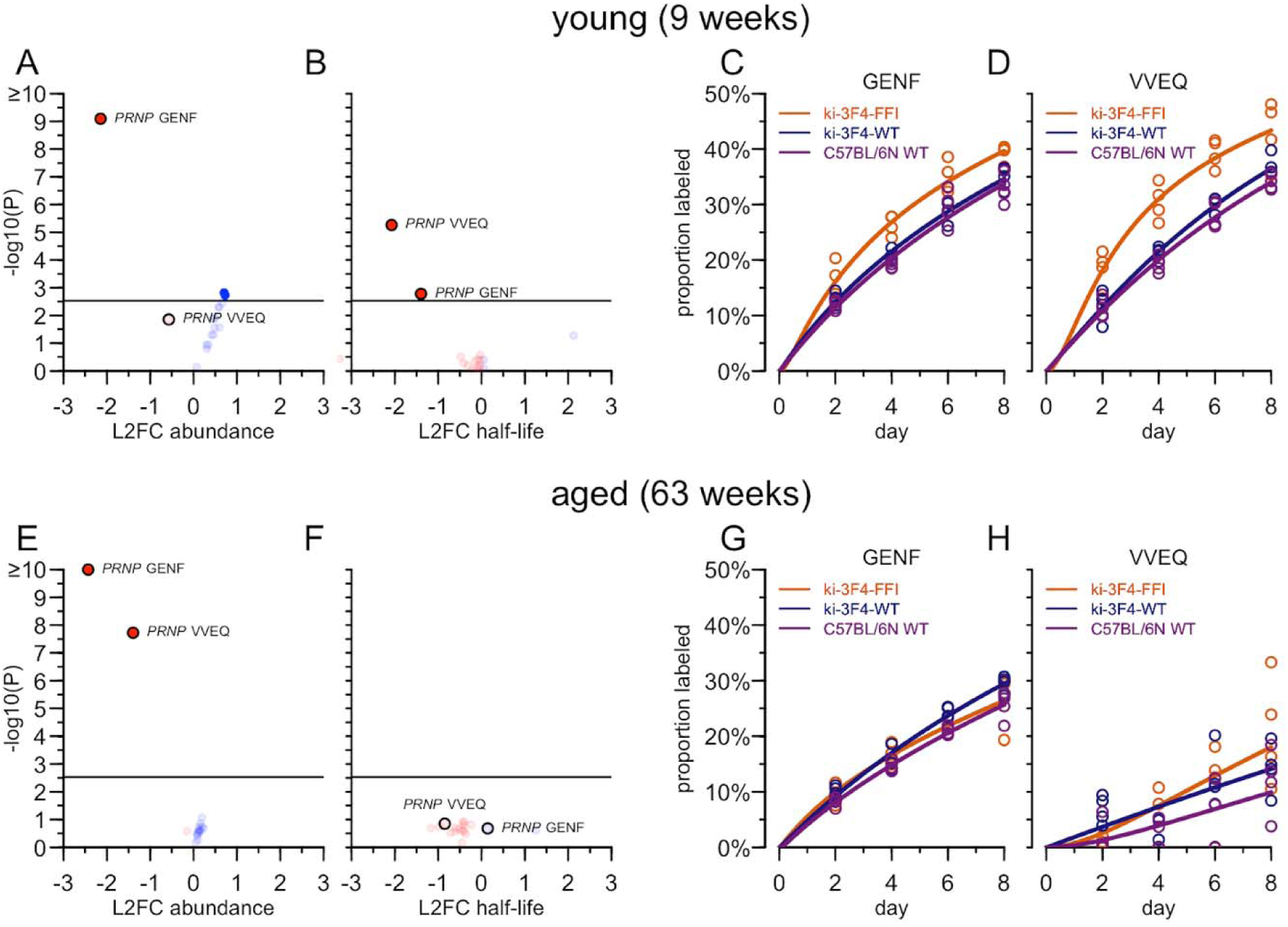
PrP turnover in mice with the equivalent of the D178N pathogenic variant by targeted mass spectrometry. **A)** Peptide abundance across all young animals. Each point is one peptide included in this targeted MS assay. The x axis position is the log2 fold change in total intensity (light + heavy peptide) in ki-3F4-FFI animals (N=16) versus combined ki-3F4-WT and C57BL/6N animals (N=32 total). The y axis is-log10(P), with the P value from a 2-sided T-test. **B)** Half-life across all young animals. Each point is one peptide included in this targeted MS assay. A half-life is fit to the labeled chow days and proportion heavy peptide across all ki-3F4-FFI animals (N=4 per timepoint times 4 timepoints) and compared to that fit to the combined control groups (N=8 per timepoint times 4 timepoints). The x axis position is the log2 fold change in this estimated half-life. The y axis is-log10(P), with the P value of the main effect of genotype (ki-3F4-FFI vs. combined controls) in a beta regression model for proportion labeled peptide as a function of chow days. **C)** Label accumulation in the GENF peptide by genotype. Each point is one animal. The x axis position is the number of days of labeled chow, the y axis position is the proportion of the GENF peptide that is heavy lysine labeled. Solid lines represent mixture-of-rates decay model fit to the data. **D)** As in C but for the VVEQ peptide. **E-H)** As in A-D but for aged, 63-week old mice.

In aged mice, the total abundance of PrP peptides was again reduced in ki-3F4-FFI (Figure 4E and S8), confirming underexpression, however, no difference in half-life was detected (Figure 4F). The accumulation of heavy peptide over time in ki-3F4-FFI mice was indistinguishable for either peptide (Figure 4G-H).

Having observed in the PrP-lowering active ASO 6 experiment (Figure 3A) that a single-rate model visibly failed to adequately fit the data, we asked whether the same phenomenon could be replicated by mass spectrometry (Figure S9). We fit both single population exponential decay models and mixture-of-rates models to the data for both VVEQ and GENF peptides for each age group and genotype. The mixture-of-rates models achieved visibly tight fits to the data (Figure 4C,D,G,H). Comparing the quality of fits from single population versus mixture models, we found that the Akaike Information Criterion (AIC) favored the mixture-of-rates model in 9/12 cases, with the 3 exceptions all being VVEQ peptide in aged mice. On average, the mixture-of-rates models suggest across the 9 remaining cases that a fast component comprises 27.3% [95% CI: 17.5% to 37.2%] of the total PrP population with a very short half-life of 0.068 days [95% CI: 0.01 days to 0.152 days]. The slow component was estimated to be on average 72.7% [95% CI: 62.8% to 82.5%] of the total PrP population with an average half-life of 6.80 days [95% CI: 5.26 days to 8.34 days].

Modeling indicated that the main effect of the ki-3F4-FFI genotype on proportion labeled peptide was highly significant (shown in Figure 4B) while the interaction term between genotype and days of labeled chow was non-significant, implying no additional effect of ki-3F4-FFI beyond the first 2 days.

## Discussion

Our data indicate that PrP’s half-life is between 4.8 to 6.4 days in brain parenchyma, regardless of PrP expression level, regardless of human or mouse PrP amino acid sequence, and regardless of prion infection status, when modeled as a homogenous population. Our estimate is in agreement with prior studies using mass spectrometry on the brains of mice fed isotopically labeled chow [15,16]. A conflicting report from Safar et al determined a half-life of 0.75 days [14]. One explanation for this discrepancy is that study was performed in a transgenic Tet-off mouse line with PrP under a foreign promoter [36]. That mouse line apparently exhibited a PrP expression pattern different from endogenous PrP, as evidenced by the perinatal lethality observed when PrP was not suppressed with doxycycline treatment during pregnancy [36]. This might account for the different result. A second point is that study used antibody D13, which recognizes an N-terminal epitope upstream of the alpha cleavage site in PrP, whereas our mass spectrometry and ELISA approaches all relied on C-terminal epitopes and peptides (Figure 2A). C1, the C-terminal PrP fragment resulting from alpha cleavage [37], is reported to nearly as abundant as full-length PrP in the brain [38]. If C1 turns over more slowly than full-length PrP, this might also contribute to the discrepancy between studies. In our studies of human CSF PrP in prion disease and of rat CSF and brain PrP upon ASO treatment, however, we have found that measurement of PrP by N- versus C-terminal peptides yields the same results [17,23]. A third consideration is that the mouse model used by Safar et al also overexpresses PrP, and the animals in that study were infected with prions, but our data argue that neither of these factors is likely to explain the lower half-life estimate. In prion-infected mice dosed at 105 dpi and harvested from 106 to 133 dpi, we measured a half-life similar to that in uninfected mice, suggesting that the reported [39] downregulation of normal PrP^C^ occurring by 120 dpi, if true, might not arise from accelerated turnover.

We find no evidence that CSF PrP lags brain PrP. In a previous study, where we analyzed only rat cerebrum (cortex and subcortex) we found that at 4 weeks post-dose, each 1% PrP lowering in CSF corresponded to 1.4% lowering in cerebrum [17]. Our present data, analyzing both cerebrum and cerebellum, indicate that this discrepancy might arise not from a lag in CSF PrP response to brain PrP concentration changes, but rather, from weaker target engagement in the cerebellum, which is also in contact with CSF.

We report that PrP in colon, while lower than brain, is quantifiable by ELISA. This is consistent with colon being the only tissue besides CNS and PNS to be affected in prion disease: patients with C-terminal truncating mutations that remove PrP’s GPI anchor and cause a gain of function through change in localization often experience chronic diarrhea misdiagnosed as inflammatory bowel disease for decades before the onset of peripheral neuropathy and then dementia [40,41]. Unlike those other peripheral tissues with lower PrP expression, colon is highly innervated; the relatively high PrP concentration there might reflect enteric nervous system expression. While colon biopsy is not practical in human trials, colon PrP quantification should serve as a useful proxy for peripheral target engagement in animal studies of systemic PrP-lowering therapeutics, particularly for small molecule or antibody programs where early proof of mechanism may help to motivate the engineering effort required to cross the blood-brain barrier in sufficient quantity to engage PrP. In mice fed isotopically labeled chow, heavy PrP peptide accumulates about twice as fast in colon as it does in brain. By examining the ratio of single heavy versus double heavy among double-lysine proteins, we determine that dietary heavy amino acid availability is more rapid in colon than brain, possibly due to direct incorporation of dietary amino acids bypassing the bloodstream. After accounting for this amino acid availability difference, the estimated half-life of PrP in colon appears just slightly faster than that in brain. One possible explanation may be that cell division in the colon contributes to PrP turnover.

The D178N variant in PrP, which causes highly penetrant genetic prion disease in humans [42], results in diminished PrP concentration in CSF even in asymptomatic at-risk patients [19,20], and is also underexpressed in mouse brain and in cultured cells [28,35]. We therefore utilized the ki-3F4-FFI mouse model, which harbors the equivalent of this variant, to examine the effect of this amino acid substitution on PrP turnover. We confirmed underexpression of PrP in brain parenchyma, but observed accelerated PrP turnover only in young mice, where the half-life of mutant PrP was estimated at 1.8 – 2.4 days. In aged ki-3F4-FFI mice, PrP turnover was indistinguishable from control animals. This may provide partial support for the hypothesis that the mid-to late-life onset of genetic prion disease could be triggered by progressive malfunction in protein quality control that, in youth, helps to clear the mutant protein. Our data are not fully consistent with this hypothesis, however, because we still observed diminished steady state levels of mutant PrP even in aged animals. Regardless, our data suggest that mutant PrP will turn over faster, if anything, than wild-type PrP, and thus that the time to effect for a PrP-lowering drug will be shorter, if anything, in patients with genetic prion disease.

The diminution of PrP after ASO treatment is not perfectly modeled by a single exponential decay curve. Compared to the theoretical model, we observe deeper target engagement in the first few days after dosing than should be possible given that even the RNA has not been fully suppressed and less than one PrP protein half-life has passed; conversely, we observe less deep target engagement after 2-3 weeks than should be expected. Likewise, in mice with the equivalent of the D178N variant, which is underexpressed, all of the increase in heavy peptide accumulation after exposure to isotopically labeled chow accrued within the first 2 days. One interpretation is that there are multiple populations of PrP that degrade according to different kinetics: for example, PrP might have different half-lives on different cell types, in different subcellular compartments, or in different multimeric states or glycoforms. Our observations justify this interpretation because they match the predicted theoretical behavior of a ground truth heterogeneous population with fast and slow subpopulations modeled with a homogenous rate. In cases where the single rate models visibly fail to fit, model selection criteria such as AIC greatly preferred the mixture-of-rates model despite the penalty for increased model complexity. The half-life of the slow component in the heterogenous population is always going to be longer than a single rate estimate, and may thus point towards a need for later timepoints in some experiments. Nonetheless, our mixture-of-rates models generally suggest that the majority of PrP in the brain belongs to a “slow turnover” population with a half-life of ∼5-9 days, not far from the best half-life estimate obtained for a single population model.

Our study has limitations. Our use of ^13^C_6_ lysine-labeled chow limited us to terminal lysine peptides, which are located in PrP’s C-terminal domain; the two N-terminal peptides measured previously [23] have terminal arginines and could not be used here, precluding interrogation of PrP cleavage products. PrP is also variably N-glycosylated, being di-, mono-, or un-glycosylated, and we were unable to examine whether these species of PrP turn over at different rates. We also have not yet tested the kinetics of very deep PrP knockdown, below 50% residual. We lack a method for interrogating the half-life of PrP protein at the single cell level, so we do not know whether the rates may differ on distinct cell types. Findings in the ki-3F4-FFI mouse model we utilized here could be confounded by prion disease-related changes, particularly in the aged animals. In our colony we did not observe premature mortality in this mouse line and were unable to identify any unambiguous histopathologic or biomarker signs of prion disease through >100 weeks of age [43], however, in other laboratories this mouse line has been reported to spontaneously produce transmissible prions and to develop a variety of behavioral abnormalities [28,44]. Perhaps the most important limitation of this study is our reliance on animal models: while we modeled human PrP in transgenic mice, we have not yet studied the half-life of PrP in humans.

Overall, our data confirm that PrP’s half-life is one rate-limiting step in the time to effect of therapeutics that act by inhibiting PrP synthesis. Ideally, drugs targeting PrP production could one day be combined with drugs increasing PrP catabolism or blocking its conversion to a misfolded form; pharmacologic proof-of-concept for such approaches in vivo is lacking but more research is merited. In the meantime, some sponsors might choose to enrich for patients with relatively high functional scores or relatively slower-progressing genotypes in order to see the strongest effect in trials. Accelerating diagnosis through better neurologist awareness, rapid referral, and shortened turnaround times for diagnostic tests will be critical for reaching patients early enough to achieve sufficient target engagement while they still possess quality of life.

## Methods

### Study Approval

All animal experiments were approved by Institutional Animal Care and Use Committees (IACUC) at the Broad Institute (protocol 0162-05-17), Weissman Hood Institute (protocol 2024-AG-77) or Ionis Pharmaceuticals (protocol 2021-1176).

### Animals

All mice were of a pure C57BL/6N or mixed 6N/6J background. Details of ages and genotypes are provided in respective sections below. We utilized the ZH3 line of PrP knockout mice, the Tg25109 line of HuPrP 129M humanized mice, and the Tga20 line of PrP-overexpressing mice. Genotyping was performed by Transnetyx. Suggested primers for Tg25109 have been reported [26]. The Ki817 huPrP 129V mouse line is described here for the first time; see methods below and Figure S4. Tga20 mice were developed by Fischer et al [27]; the Tga20 transgene array was localized to the *Ptcra* gene locus by Taconic/Cergentis using targeted locus amplification (TLA) [45]; for details see Figure S5.

### Generation of human PrP knock-in mice

Humanized KI *PRNP* mice (ki817) were generated by Taconic using ES cell targeting using CRISPR/Cas9. The targeting vector was constructed using human BAC RP11-61G12 for the human *PRNP* (129V) sequence and the mouse BAC RP23-369F12 for the homology arms. This targeting vector, a puromycin resistance cassette plasmid, and a plasmid containing Cas9 and guide RNAs against desired cut sites in the mouse genome were co-transfected into C57BL/6NTac embryonic stem cells. These were then selected, incorporated into embryos, and implanted to yield founders. The full targeting strategy is provided as a PDF in this study’s online data repository. The gRNA sequences used to target the mouse genomic region to be replaced with the human sequence were GGTCTGCTGATCCGACAACG and TAGAAGCTATGATGAACACC. The exact coordinates of human sequence incorporated into the mouse span from 306 bp upstream of the transcription start site to 1 bp downstream of the transcription end site (Figure S5).

### Isotopic labeling of mice

Where necessary, prior to study start, mice were consolidated into cages of a single genotype. Mice at Broad (wild-type, ZH3/+, Tga20 heterozygous, Tg25109 heterozygous on a ZH3/ZH3 background) were between ages of 21-23 weeks old. Mice at Weissman Hood Institute (ki-3F4-FFI, ki-3F4-WT, and C57BL/6N) were 59-60 days old (∼9 weeks, young cohort) or 429-439 days old (∼63 weeks, aged cohort) at the start of isotopic diet. Mice first received fed Mouse Express (unlabeled) irradiated mouse feed (Cambridge Isotope Laboratories, USA) for 7 days at libitum (Broad) or 14 days (Weissman Hood Institute) to acclimate to the new diet. The next day, mice were switched to Mouse Express L-Lysine (^13^C_6_, 99%) irradiated mouse feed (Cambridge Isotope Laboratories, USA) ad libitum and were sacrificed after 8 days (Broad) or 2, 4, 6, or 8 days (Weissman Hood Institute). Animals had standard water access and no alternate food source was made available during the study. In the pilot study at Broad, one animal per genotype was set aside (not fed isotopic chow) as a representative control. In the pilot study, the amount of chow given each day was weighed prior to feeding and the remaining chow was weighed each day to estimate the amount of chow being eaten per animal per day. Animals were weighed on day 1 of unlabeled chow, day 1 of isotopically labeled chow and were harvested 24 hours after the last labeled chow refresh.

### Organ harvest

Animals were weighed directly before harvesting. Animals were euthanized via CO_2_ asphyxiation and disarticulation of the skull and cervical vertebrae was utilized as a secondary measure. Left and right brain hemispheres were collected by hemi-secting the full intact brain with a scalpel on ice, hemispheres were collected into separate tubes and flash frozen on dry ice. Left and right sciatic nerves were collected in length from the proximal hip to distal knee and flash frozen on dry ice. Colon sections were harvested in length from the caudal end of the ascending colon to the middle of the transverse colon and flash frozen on dry ice. All samples were stored at-80C and sent for LC-MS in the form of frozen, intact tissue.

### Immunoblots

All samples were homogenized in cold PBS with 0.2% CHAPS. All organs were homogenized at 10% wt/vol, except for sciatic nerve, which was 5% wt/vol due to limited sample mass, and blood and plasma, which were not homogenized. Each sample was then diluted 4-fold into RIPA buffer (20 mM Tris-HCl pH 8.0, 140 mM NaCl, 1 mM EDTA, 1% Triton X-100, 0.1% SDS) with protease inhibitors (MilliporeSigma 4693159001), vortexed for 1 minute, centrifuged 14,000 x G for 10 minutes at 4°C and then supernatants were diluted a further 4-fold into 4X LDS (ThermoFisher NP0007) + 50 mM TCEP (ThermoFisher 77720) and incubated 5 minutes at 95°C. 10 µL of this solution was then loaded per lane and run on a SDS-PAGE gel (4-12% Bis-tris NuPAGE No. NP0323BOX) in MES buffer for 40 minutes at 180V. Proteins were either Coomassie stained or transferred to iBlot using 20V for 7 minutes, cooled for 3-5 minutes, blocked with TBS blocking buffer (Licor No. 927-60001) for 1 hour at room temperature. 6D11 anti-PrP primary antibody (Biolegend no. 808003) was diluted 1:1000 in TBS blocking buffer with 0.2% Tween and incubated overnight at 4°C. Blots were then rinsed 4x with TBST (25 mM Tris, 0.15 M NaCl, 0.05% Tween-20 at pH 7.5; 5 minutes each with rocking), and goat anti-mouse IRDye 800CW (Licor No. 926-32210) was diluted 1:10,000 in TBS blocking buffer with 0.2% Tween, incubated 1 hour at room temperature with rocking, and washed 4x with TBST (5 minutes each with rocking). Blots were imaged at 800 and 700 nm on a Licor Odyssey CLx Infrared Imaging System.

### Tissue homogenization for ELISA

Each brain hemisphere was added to a 7 mL Precellys tube with pre-loaded zirconium oxide beads (Precellys, Bertin, USA) and homogenized in ice cold 0.02% CHAPS in 1x PBS with protease inhibitors (1 Roche cOmplete tablet 4693159001, Millipore Sigma, USA, per 10 mL of buffer) using 3x 40 seconds pulses at 6,000 rpm in the Bertin Technologies Precellys Evolution Touch Homogenizer (Bertin, USA). The final protocol for colon homogenization (after optimization, see Figure S1) used 2 mL Precellys tubes and 5x 40 second pulses at 8,000 rpm. Homogenates were aliquoted into multiple 40 µL aliquots for protein analysis and 1 mL aliquots as a backup stock, flash frozen on dry ice and stored at - 80°C until further analysis.

### PrP ELISA

PrP concentration in the rat and murine brain hemispheres was measured using a previously published PrP ELISA [17]. The capture antibody, EP1802Y (ab52604, Abcam, USA), is incubated in a clear 96-well plate overnight at 4C. After blocking and sample incubation, biotinylated 8H4 antibody (ab61409, Abcam, USA) is used for detection with streptavidin-HRP (Pierce High Sensitivity, 21130, Thermo Fisher Scientific, USA) and TMB substrate (7004P4, Cell Signaling Technology, USA). Brain homogenates and QCs were diluted to 1:200 final concentration for the assay; the final protocol for colon homogenization (after optimization in Figure S1) utilizes a 1:100 final dilution. Recombinant mouse PrP, (MoPrP23-231) prepared as described [46,47] was used for the standard curve. Average residual PrP was calculated by dividing the amount of residual PrP in each treated brain by the mean concentration of residual PrP in the vehicle and/or no dose control brains from the same study and time point. For the colon data in Figure 1D-E, the detection mAb concentration was doubled (0.50 µg/mL instead of 0.25 µg/mL), but with further assay development (Figure S1) we were able to revert to the original 0.25 µg/mL concentration and maintain the ∼5-fold margin above LLQ.

### Prnp qPCR

The qPCR procedure has been described previously [1]. *Prnp* RNA levels were normalized first to housekeeping gene *Ppia* then to the mean of PBS-treated controls. Primers are as follows. Prnp forward: TCAGTCATCATGGCGAACCTT, reverse: AGGCCGACATCAGTCCACAT, probe: CTACTGGCTGCTGGCCCTCTTTGTGACX. Ppia forward: TCGCCGCTTGCTGCA, reverse: ATCGGCCGTGATGTCGA, probe: CCATGGTCAACCCCACCGTGTTCX.

### Mouse inoculation

Mouse inoculation has been described previously [1]. Animals were freehand inoculated halfway between the right ear and right eye, approximately 1 mm right of the midline, with 30 µL of a 1% (wt/vol) brain homogenate from terminally sick RML prion-infected mice.

### Mouse intracerebroventricular (ICV) injection

Mouse ICV was similar to that described previously [1]. ASOs were diluted to 500 µg in a 10 µL dose volume in dPBS (Gibco 14190) and administered into CSF by bolus ICV injection in stereotaxis (ASI Instruments, SAS-4100).

Positioning utilized 18° ear bars in ear canals and incisors in the mouse adapter tooth bar, adjusted to −8 mm. A 1 cm incision was made and the periosteum was scrubbed with sterile cotton-tipped applicators in order to reveal bregma. Drug was administered in Hamilton syringes (VWR 60376-172) fitted with 22-gauge Huber needles (VWR 82010-236). The needle was aligned to bregma and then moved 0.3 mm anterior, 1.0 mm right, and then downward either 3.0 mm past where the bevel disappeared into the skull or 3.1 mm past where the tip of the needle first touched the skull. Liquid was ejected over 10 seconds and the needle withdrawn 3 minutes later while applying downward pressure on the skull with a cotton-tipped applicator.

Incisions were closed with a horizontal mattress stitch (Ethicon 661H suture). Animals recovered from the anesthesia in their home cages on a warming pad.

### Rat ICV injection

The rat ICV injection was as described previously [17]. The procedure is similar to that for mice (see above) except that it utilizes 27° atraumatic ear bars (ASI Instruments, EB-927), with coordinates: riser –6 mm, 1 mm caudal, 1.5 mm right, 3.7 mm from the surface of the brain into the lateral ventricle. A bore hole was first drilled using a sterile 1 mm × 33 mm drill bit (McMaster Carr, 5058N51) in a hanging-style handpiece (McMaster Carr, 4454A14) held in a stereotactic handpiece holder (ASI Instruments, DH-1000). Injection volume was 30 µL in a gastight 1710 small RN syringe (Hamilton 81030). Incision closure utilized 5-O monofilament suture (Ethilon 661G-RL).

### Targeted mass spectrometry

Peptides terminating in lysine were nominated based on prior mass spectrometry work [23]. Single peptide quantification for VVEQMCVTQYQK (Figure 2A) was performed at Charles River Labs (Worcester, MA). Serial dilution experiments determined an LLQ of 0.111 ng/mL (ng of peptide per mL of homogenate), corresponding to 30.6 fmol/mg (fmol of peptide per mg of total protein). Multiplex quantification of VVEQMCVTQYQK, GENFTETDVK, and the control peptides from other proteins (Figure 2D-F) was performed at IQ Proteomics (Framingham, MA). Details of the LCMS methods are provided in the supplementary material.

### Labeled peptide accumulation models

Free lysine in plasma was assumed to follow the fit described by Fornasiero [16]: H = 1-0.503*(exp(-t*ln(2)*0.799))-0.503*exp(-t*ln(2)/39.423), where H means the proportion of free lysine that is ^13^C_6_ labeled, and exp(x) signifies e^x^. Using this formula we calculated the percent labeled at every timepoint from 0 to 8 days using increments of dt = 0.01 days. We then calculated the accumulation of ^13^C_6_ lysine label in peptides using numerical integration as follows. For parameter lambda (λ), defined as λ = ln(2)/t_1/2_ the turnover of protein in an arbitrarily small unit of time dt is λ*dt. For a 5-day half-life protein, for example, λ = ln(2)/5 = 0.14, so that every 0.01 days, 0.01*ln(2)/5 = 0.0014 (expressed as a proportion) or 0.14% of protein is catabolized, and 0.14% of the original amount is produced to replace it. The protein begins 100% unlabeled, and label accumulates as unlabeled protein is catabolized and a greater and greater proportion of the nascent protein is ^13^C_6_ labeled. The uniroot function in R was used to perform the inverse operation — estimating a half-life from the observed proportion labeled, by minimizing the residuals between this numerical integration model and the actual data over 1 or more observed timepoints. To assess genotypic differences in label accumulation, we fit a beta regression model where proportion of peptide labeled is a function of days of isotopically labeled chow and genotypic group, with an interaction term: betareg(prop_labeled ∼ chow_days * grp). We chose a beta regression model because the proportion of peptide that is labeled is inherently constrained to be between 0 and 1 inclusive, which this model reflects. The main effect for the genotype group is interpreted to signify the difference in peptide labeling equally affecting all timepoints, while the interaction term is interpreted to signify the difference in label accumulation over the timepoints.

### Free amino acid availability estimation

To empirically estimate the availability of dietary labeled lysine in brain and colon, we used an approach described by Beynon and colleagues [29,30]. If the proportion of free lysine available for protein synthesis that is heavy labeled, or relative isotope abundance (RIA) is p, then among peptides with two lysines, the proportion (1-p)^2 will be light-light (LL), 2p(1-p) will be light-heavy (LH), and p^2 will be heavy-heavy (HH). All light-heavy and heavy-heavy peptides represent the result of nascent protein synthesis during the labeling period, and thus, availability of heavy lysine can be estimated without knowledge of protein half-life. The ratio of LH:HH is 2p(1-p)/(p^2). Solving this equation for p reveals that p = 2/(LH/HH + 2). In order to obtain quantitative information on double-lysine peptides, we performed a limited trypsin digest with a 1:100 enzyme:protein ratio and a digestion time of 1.5 hours (other parameters were as described in Supplemental Methods > Targeted mass spectrometry at IQ Proteomics). Digested samples were analyzed by data independent acquisition (DIA) on an Orbitrap Astral instrument (30 min gradient), and the results were searched using Spectronaut. For double lysine peptides with quantifiable LH and HH signal, we calculated point estimtes of RIA = 2/((LH/HH) + 2). These were then grouped by tissue and by animal or by intensity bin. In order to adjust the plasma free lysine model for the higher RIA in colon, we simply multiplied the original free lysine formula by the ratio of colon RIA to brain RIA (1.67). This resulted in an estimated heavy lysine availability in colon of 93.0% by day 8.

### Exponential decay model

All residual *Prnp/PRNP* RNA and PrP protein measurements were normalized, respectively, to the mean RNA and protein measurements in the untreated animals across all timepoints so that all measurements are on a scale from 0% (complete knockdown) to 100% (normal levels). Observed *Prnp/PRNP* RNA measurements in brain were linearly interpolated using the approx function in R to yield point estimates of residual RNA at units of dt = 0.01 days. Similar to the approach described above, the residual protein (P) at any given timepoint was computed numerically as a function of RNA (R) and the exponential decay parameter λ = ln(2)/t_½_ value. Catabolism of protein is proportional to the amount of protein in the previous time increment, while synthesis of protein is proportional to RNA in the previous time increment. So, at any time t, the protein catabolized is P_t-1_ * λ * dt, and the protein synthesized is R_t-1_ * λ * dt. Thus, the change in protein dP = R_t-1_ * λ * dt - P_t-1_ * λ * dt. The amount of protein at time t is P_t_ = P_t-1_ + dP. With this function in hand, another function was written to calculate the residuals of the actual data as compared to this model. Then, the nls.lm function in R was used to determine the λ value that minimizes those residuals, this fitting the model. A time increment of dt=0.01 and a starting guess of λ = 0.14, corresponding to a half-life of 5 days, were used in fitting the model.

### Mixture-of-rates models

To test whether protein pools exhibit heterogeneous turnover kinetics, we fit two-component mixture models to both isotopic labeling and residual decay data. For isotopic labeling, the mixture model assumes the peptide pool comprises two subpopulations with distinct half-lives (t_½,1_ and t_½,2_) and proportions (π_1_ and π_2_ where π_1_ + π_2_ = 1). Label accumulation in each subpopulation was computed independently using the numerical integration approach described above, and the total proportion labeled was calculated as the weighted average π_1_·L_1_(t) + π_2_·L_2_(t), where L_1_(t) and L_2_(t) represent label accumulation in the fast and slow pools, respectively. For residual decay experiments, the mixture model similarly assumes two protein pools with distinct decay rate constants (λ_1_ and λ_2_) and initial proportions (w_1_ and w_2_). Each pool’s dynamics follow dPk/dt = λk·R(t)·wk - λk·Pk(t), with total protein P(t) = P_1_(t) + P_2_(t). Mixture models were fit using the Nelder-Mead optimization algorithm with multiple starting parameter combinations to ensure robust as possible optimization. Model selection between single-rate and mixture models was performed using the Akaike Information Criterion (AIC = n·ln(RSS/n) + 2k) and Bayesian Information Criterion (BIC = n·ln(RSS/n) + k·ln(n)), where n is the number of observations, RSS is the residual sum of squares, and k is the number of model parameters. For the single-rate model, k = 2 (λ and σ) for residual decay or k = 1 (t_½_) for isotopic labeling; for the mixture model, k = 4 (λ_1_, λ_2_, w_1_, and σ) for residual decay or k = 3 (t_½,1_, t_½,2_, and π_1_) for isotopic labeling. ΔAIC <-3 was interpreted as substantial evidence favoring the mixture model. For mixture models, the effective half-life was calculated as the time at which the combined pool reaches 50% of its initial state by numerically solving Σ(w_i_·0.5^(t/t_½,i_)) = 0.5.

### Use of large language models (LLMs)

Claude Code was used to generate first drafts of source code for a subset of analyses and figure panels. These blocks of code were subsequently reviewed and validated by DAS and EVM.

### Statistics and source code

All analyses were conducted using custom scripts in R 4.4.1, and the mixture-of-rates decay models fit using Julia 1.12.4. Exponential decay and label accumulation models are described above. Differences between genotypes in Figure 2 were compared using a 2-sided T test and Bonferroni corrected for 10 tests. Differences in peptide abundance between genotypes in Figure 4 were compared using a 2-sided T-test, while differences in labeled peptide accumulation were assessed using a beta regression model (see above). All error bars or shaded areas shown are 95% confidence intervals. P values of < 0.05 were considered significant.

### Data availability

Raw data and source code sufficient to reproduce all figures and statistics in this manuscript are available at github.com/vallabhminikel/halflife.

## Author contributions (CRediT statement)

Designed the experiments: TLC, ABS, BE, CB, AGR, WSJ, AGP, SMV, NO, HTZ, EVM. Performed the experiments: TLC, JOM, AS, VL, BN, NGK, FES, ABS, BE, CB, KL, MH, NC, AGR, DEC. Supervised the research: SMV, AGP, NO, HTZ, EVM. Analyzed the data: TLC, DAS, YL, ABS, CB, NO, HTZ, EVM. Drafted the manuscript: TLC, DAS, ABS, CB, EVM. Reviewed and approved the final manuscript: all authors.

## Acknowledgements

This study was supported by the National Institutes of Health (R01 NS132022 to EVM and COBRE P20 GM152335 to AG), Prion Alliance, Ionis Pharmaceuticals, and Gate Bio. The funders had no role in study design, data collection and analysis, decision to publish, or preparation of the manuscript. HTZ and BN received salaries from Ionis Pharmaceuticals and NO received salary from Gate Bio.

## Competing interests

We have read the journal’s policy and the authors of this manuscript have the following competing interests. BE and CB are employees of IQ Proteomics. ABS is an employee of Charles River Laboratories. NO is an employee and shareholder of Gate Bio. HTZ and BN are employees and shareholders of Ionis Pharmaceuticals. EVM has received speaking fees from Abbvie, Eli Lilly, Novartis, Vertex, and Voyager; consulting fees from Alnylam, Arrowhead, Deerfield, and Regeneron; and research support from Cenos, Eli Lilly, Gate Bio, Ionis, Regeneron, and Sangamo Therapeutics. SMV acknowledges speaking fees from Abbvie, Biogen, Eli Lilly, Illumina, Ultragenyx, and Voyager; consulting fees from Alnylam, Invitae, and Regeneron; research support from Cenos, Eli Lilly, Gate Bio, Ionis, Regeneron, and Sangamo Therapeutics.

## Notes

https://github.com/vallabhminikel/halflife

### Summary of Updates

Updates in response to reviewers round 1

