## Supplement for "PrP turnover in vivo and the time to effect of prion disease therapeutics"

Corridon et al 2026

**Supplementary Figures**

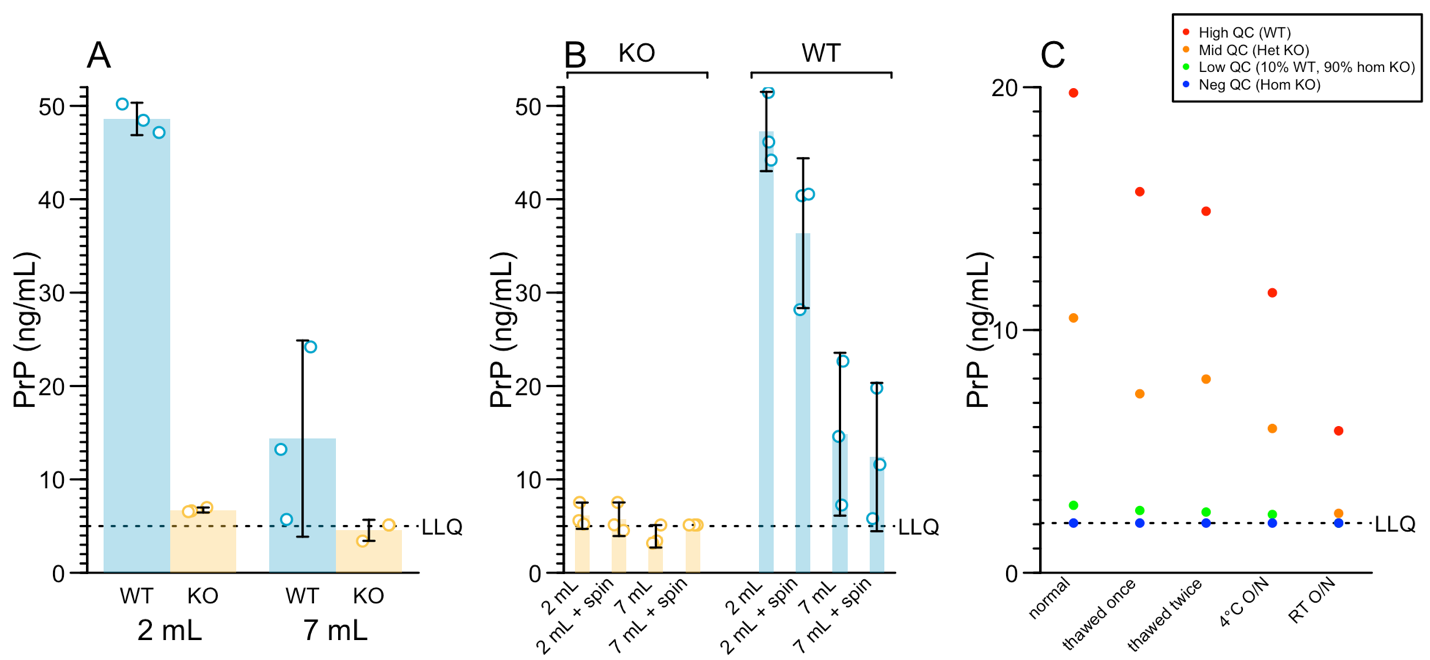

***Figure S1. Assay development for ELISA quantification of PrP in colon. A)*** *Effect of homogenization tubes used. 7 mL tubes homogenize incompletely, leading to lower detection. These data used our standard ELISA conditions with 0.25 µg/mL detection antibody and were conducted at a 1:100 dilution.* ***B)*** *Centrifugation of samples does not rescue the under-recovery of PrP when 7 mL homogenization tubes are used.* ***C)*** *Stability study. PrP in colon samples is subject to loss upon freeze/thaw, time at 4°C or time at room temperature (RT) overnight (O/N). For this stability experiment we treated the lowest standard curve point, 0.02 ng/mL, as the lower limit of quantification (LLQ).*

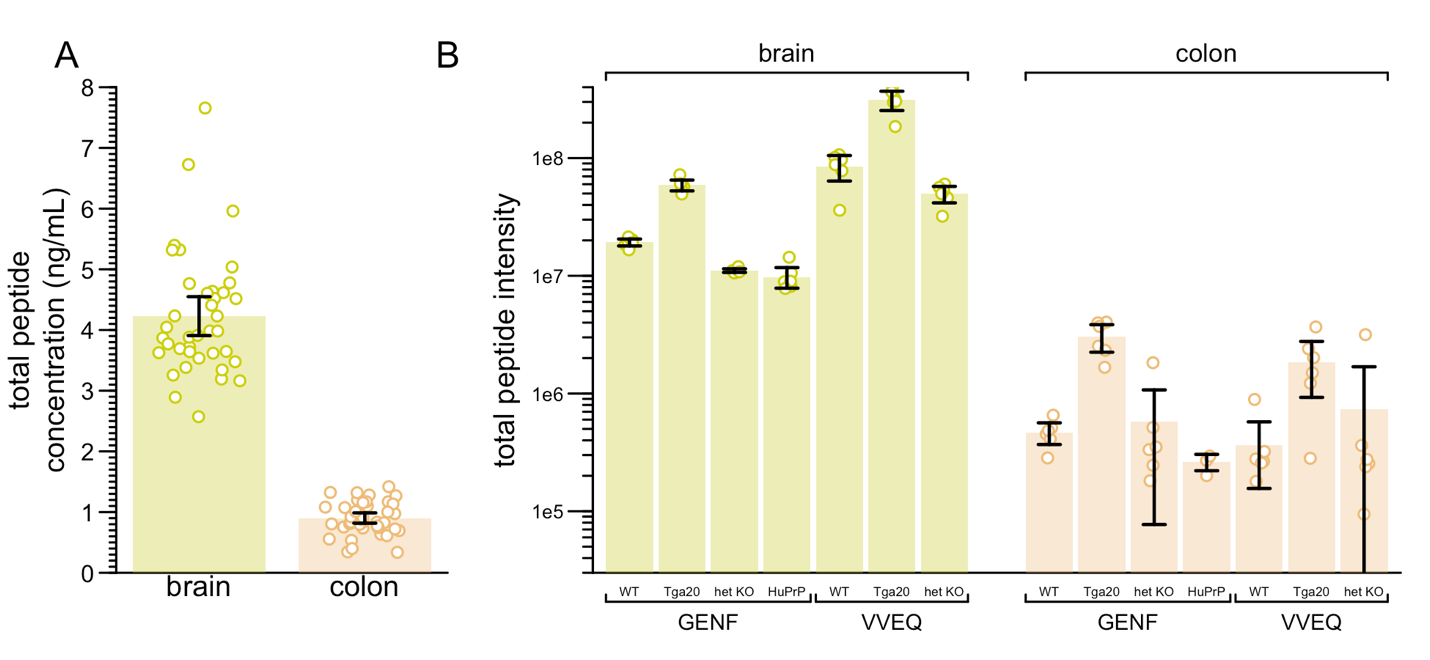

***Figure S2. Total abundance of light+heavy peptide in the mass spectrometry assays. A)*** *The Charles River Labs (CRL) assay for VVEQ. Brain averages 4.7 times higher than colon.* ***B)*** *IQ Proteomics assay for VVEQ and GENF. Brain detection is 19.3 – 231 higher than colon, at odds with our Western and ELISA analysis (Figure 1) and the CRL assay (Figure S1A). Protein in colon may have been under-recovered due to incomplete homogenization. Nonetheless, this does not affect the proportion labeled measurement used in Figure 2.*

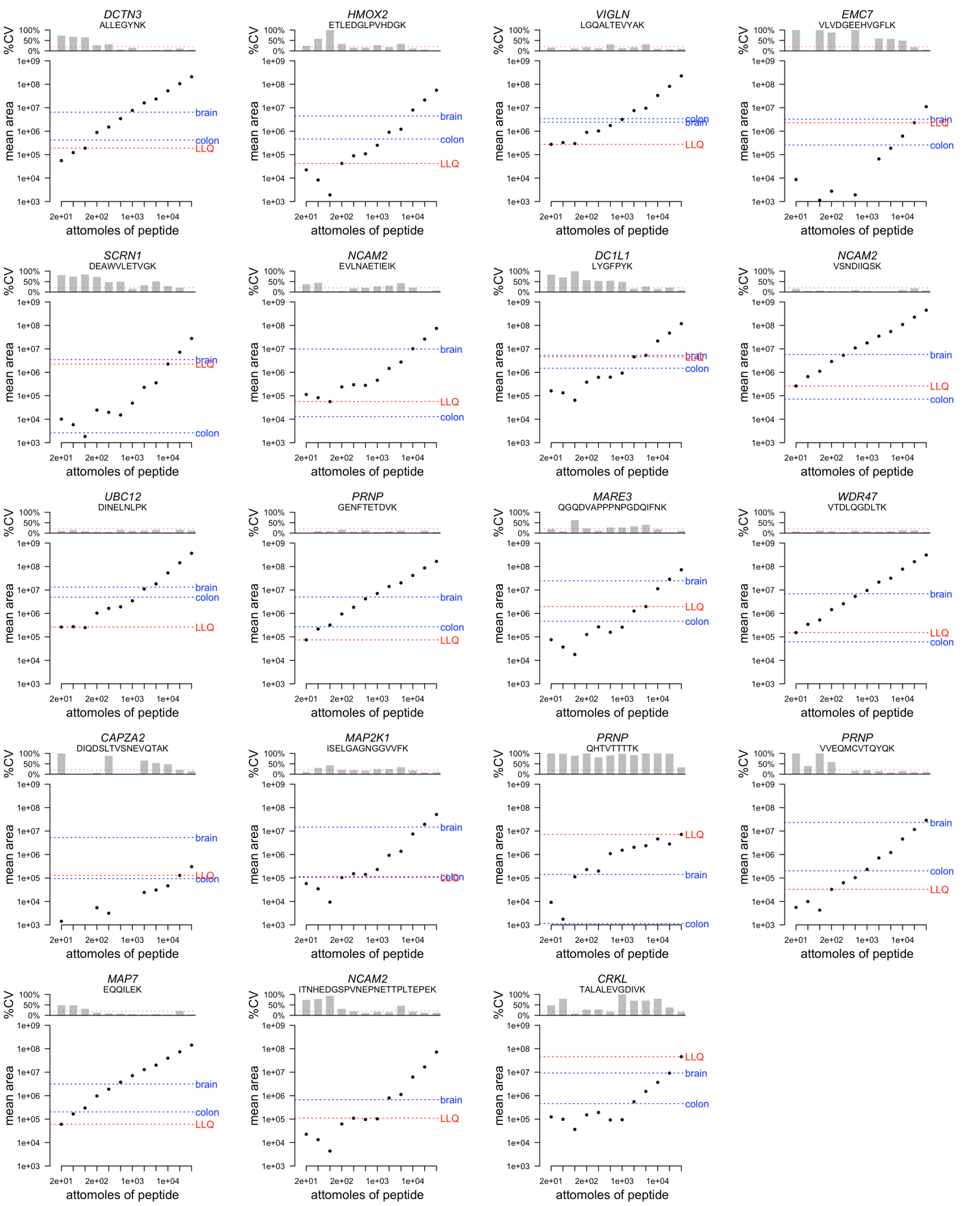

***Figure S3. Quality control and LLQ determination for the IQ Proteomics assay.*** *Each panel represents 1 of 19 peptides, whose identity and gene are shown at top. Synthetic peptides were serially diluted in triplicate. The upper section of each panel shows the coefficient of variation (%CV; y axis) among technical replicates at each dilution versus the nominal amount of peptide in amol (attomoles; (x axis shared with lower section). The lower section shows the intensity (y axis) versus nominal amount. We defined each peptide’s LLQ (red dashed line) as the most dilute concentration for which the %CV of all concentrations equal or stronger than is ≤20%. Also shown are the mean heavy peptide intensities (blue dashed lines) for wild-type mouse brain and colon at day 8. Thus, if a blue line is above the red line, this means that in wild-type mice after 8 days, the heavy peptide in that tissue is detected above the LLQ.*

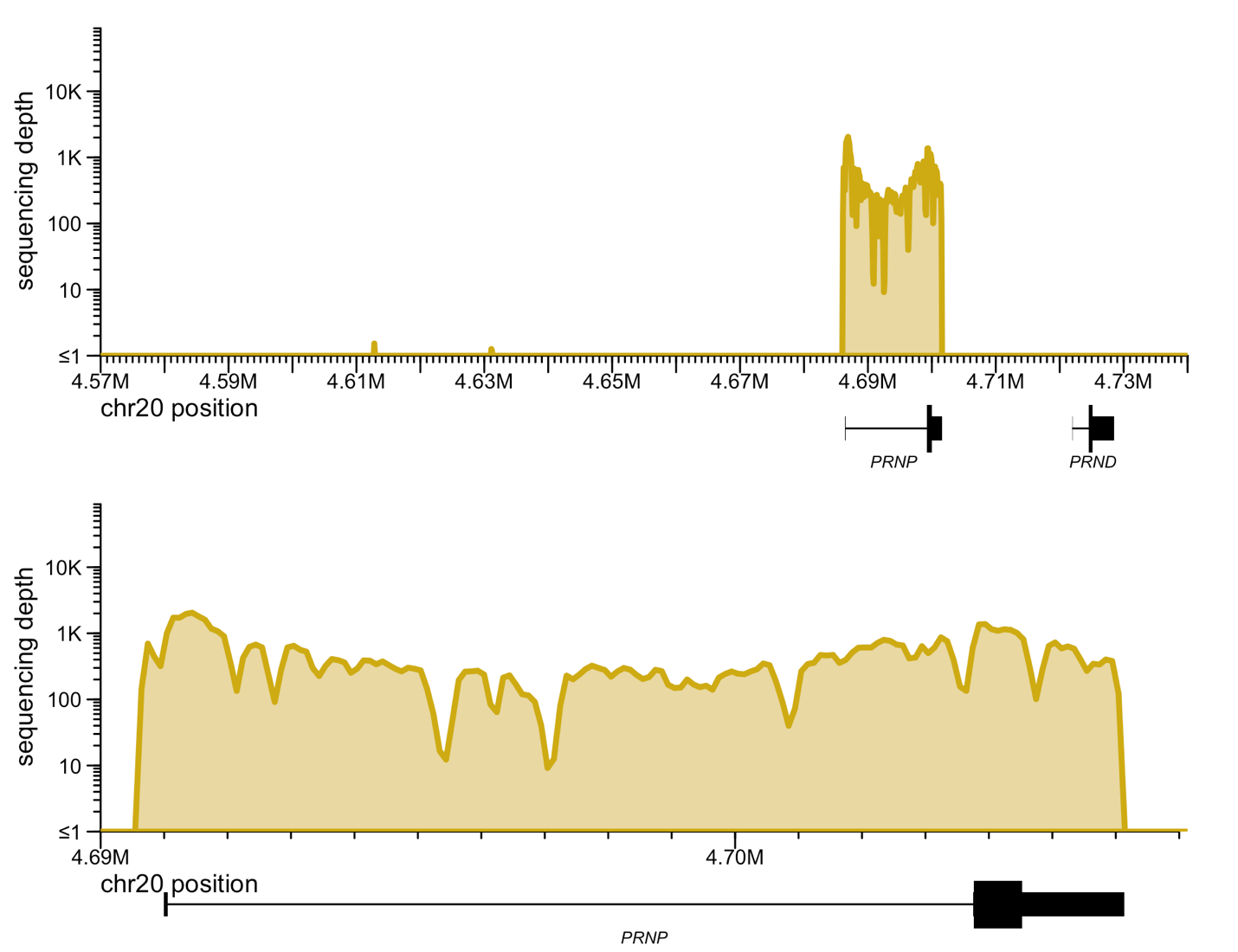

***Figure S4. Human genomic sequence in ki817 mice.*** *Genomic DNA from a ki817 mouse was subjected to targeted sequencing for 152 kb around the PRNP locus using custom baits* (1) *(Twist biosciences) and aligned to the human reference genome. GRCh38 coordinates are shown. The knock-in allele spans 306 bases upstream of the human transcription start site (TSS, located at GRCh38 chr20:4,686,456) to 1 base downstream of the human transcription end site (TES, located at GRCh38 chr20:4701588).*

*Sequencing reads demonstrating the breakpoints:*

GRCm39 chr2:131751847 (TSS +0 bp) / human GRCh38 chr20:4686151 (TSS -306bp bases upstream of the TSS):

GGCGCGGCCATTGGTGAGCATCACGCCCCGCCCCTCGCCCAGCCTAGCTCCCGCCTGCCCCGATTAAAGATGATTTTTACAGTCAATGAGCCACGTCAGGGAGCGATGGCACCCGCAGGCGGTATCAACTGATGCAAGTGTTCAAG

human GRCh38 chr20:4701589 (1 base past the TES) / GRCm39 chr2:131780358 (1 base past TES)
TGAAATTAAACGAGCGAAGATGAGCACCACGGGGTTTGTTCTCTCTCCAATGCTCCGAGTCCACTGTTTATCGCCAGGGTGGCTTGGGCTCATTTCACATCCCTGTCCCTGAGGGGCCTCGGGTCTTACCTCTGGTCCTGTCTTGT

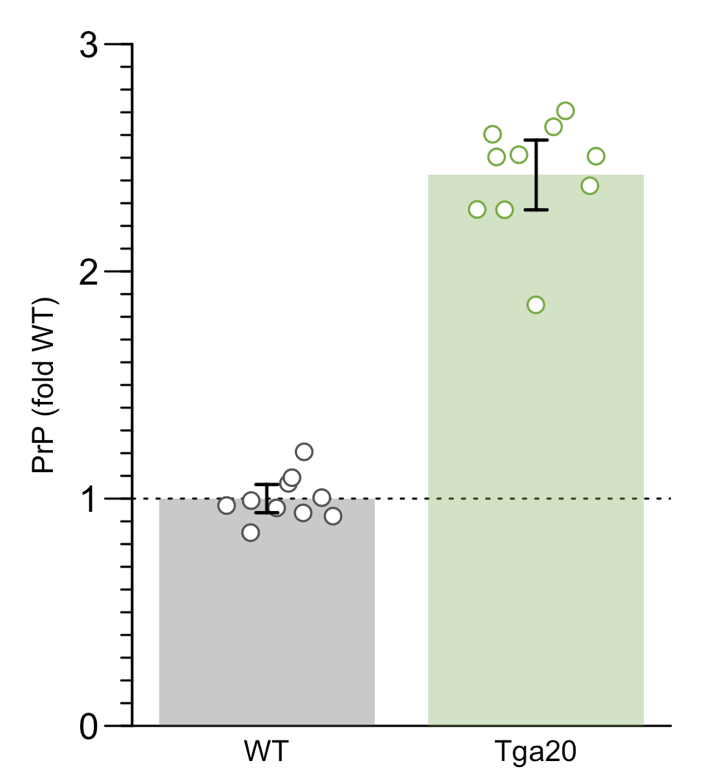

***Figure S5. PrP expression in Tga20 mice.*** *PrP ELISA on whole brain hemisphere homogenates from wild-type C57BL/6N mice or mice hemizygous for the Tga20 transgene array, on a background of endogenous PrP knockout (Tga20/0; ZH3/ZH3). The mean Tga20 value is 2.4x the wild-type result. The original report of the generation of the Tga20 line estimated that animals hemizygous for this transgene array expressed 6-7x wild-type PrP levels, by Western blot densitometry. We have shown that our ELISA provides quantitative estimates of relative PrP expression when samples are analyzed at the same dilution (2), and brain samples from all animals in this experiment were analyzed at the same 1:200 weight/vol final dilution. Transgene mapping of the Tga20 transgene array revealed integration at position (GRCm38) chr17:46,761,775-46,762,856, within intron 1 of Ptcra. The full transgene mapping report is provided in the study’s online git repository. The genomic breakpoints were identified as follows:*

5’ integration site: GRCm38 chr17:46,761,775 (tail) fused to TG (homologous to chr2:131,909,936 (head)) ATCCCAGCGCCTACACACCCAACACTTCAATCTGTAATGAAATCCTATGCCCTCGTCTAGTGTGTCTGAAGACACACTCCCGGCTCCCCCGCGTTGTCGGATCAGCAGACCGATTCTGGGCGCTGCGTCGCATCGGTGGCAGGTAAGCG

3’ integration site: TG (homologous to chr2:131,903,786 (head)) fused to GRCm38 chr17:46,762,856 (head) with 3 inserted bases CTTGTTGGAAGAAGTGTGTAATTGGGGGTGAGCTTTGAAGTTTCAAATGCTTAAGCCAGGCCCAGTATCACCCTCTCTGCCTTTTTGCTGCCTTTGGATCCGTCGCTTTGGAAATAATCTTTCTTTTTTTTAAGATTTATTTTATGTAT

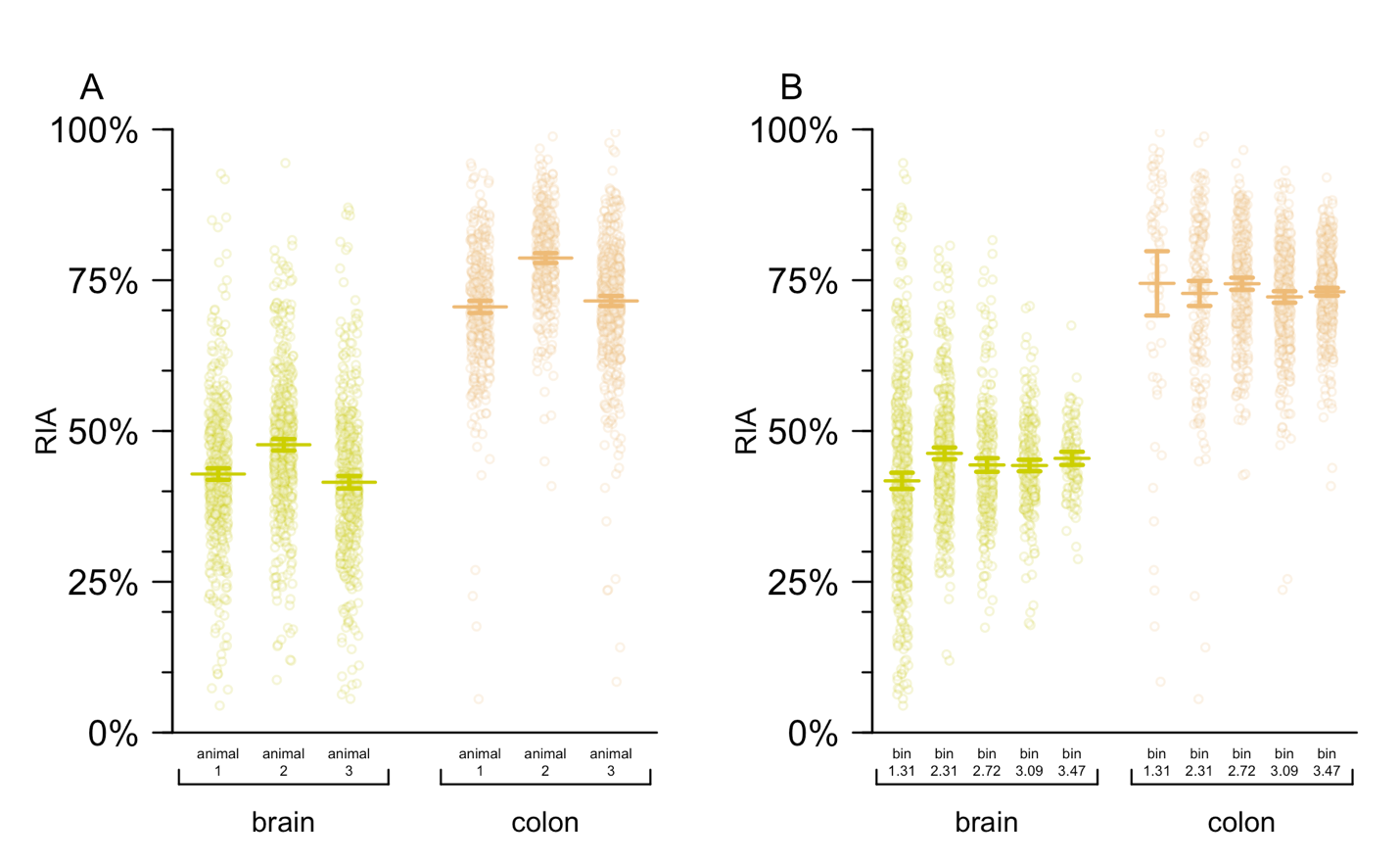

***Figure S6. Empirical determination of heavy lysine availability in brain and colon.*** *The empirically determined relative isotope abundance (RIA) calculated from N=833 light-heavy to heavy-heavy double-lysine peptides from N=596 proteins in paired brain and colon samples from N=3 wild-type C57BL/6N animals after 8 days of labeled chow.* ***A)*** *Point estimates of RIA grouped by animal and tissue. Each point is a peptide. Segments and error bars represent 95% confidence intervals.* ***B)*** *Point estimates of RIA grouped by log10 bins of intensity (peptide abundance) and tissue. Each point is a peptide-animal tuple. Segments and error bars represent 95% confidence intervals. The source data, including exact number of peptides per bin, are provided in the Supplementary Data.*

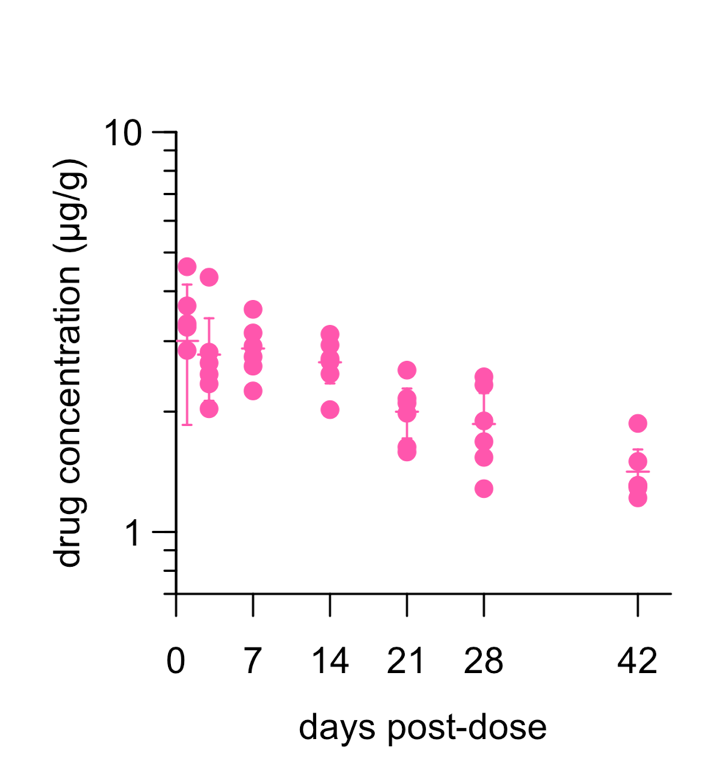

***Figure S7. Pharmacokinetic (PK) parameters of ASO N in ki817 mice.*** *Drug concentration (µg/g, y axis) versus days post-dose (x axis). Points represent individual animals (the same whole hemispheres used for qPCR), line segments represent means, and error bars represent 95% confidence intervals.*

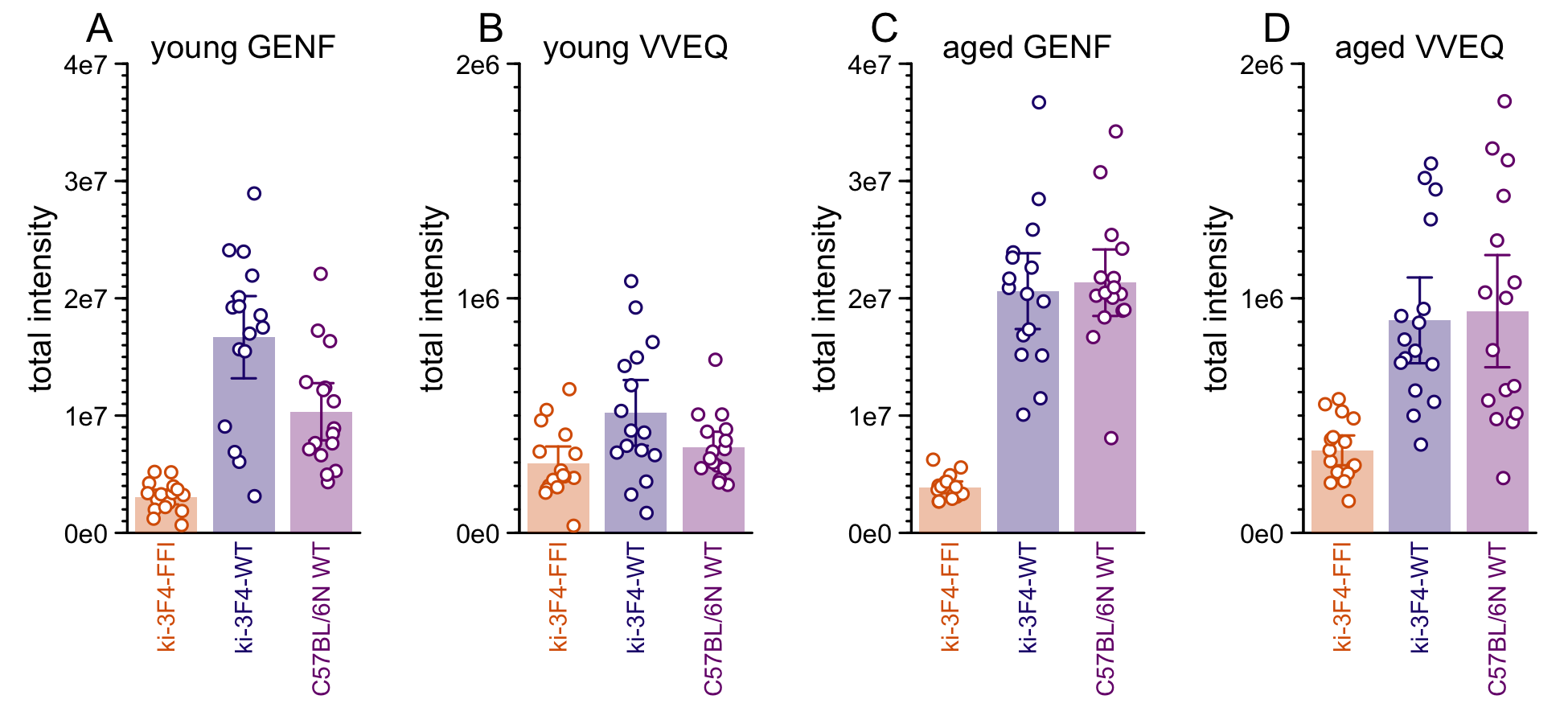

***Figure S8. PrP peptide abundance by genotype and age in ki-3F4-FFI mice and controls.*** *Shown on the y axis are total intensity (heavy + light peptide), grouped by genotype on the x axis; each point is one animal, bars represent means and error bars represent 95% confidence intervals. Age of animals and PrP peptide being monitored are indicated at top of each panel.*

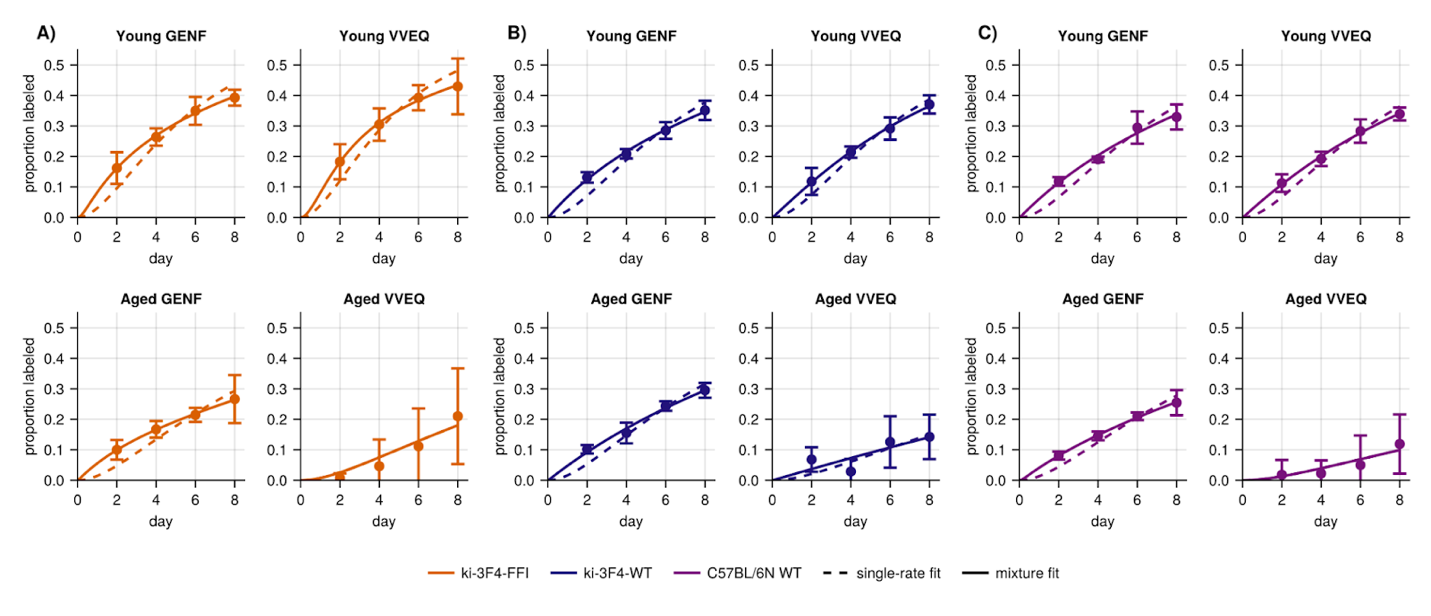

***Figure S9. Comparison of single-rate and mixture-of-rates decay models.*** *For each peptide-age combination the genotypes A) ki-3F4-FFI B) ki-3F4-WT and C) C57BL/6N WT have the label accumulation plotted. Single-rate (dashed line) and mixture-of-rates (solid line) models are shown.*

**Supplementary Methods**

***Targeted mass spectrometry at IQ Proteomics.***

*Lysis performed at IQ Proteomics.* 1mL of lysis buffer containing 8M Urea, 100 mM EPPS pH 8.5, and 1X HALT protease/phosphatase inhibitor (Pierce) was added to each sample. For complete lysis and shearing of genomic DNA, ceramic beads were added to each sample, and the samples were subjected to bead beating (Precellys Evolution) using four cycles at 5800 RPM for 30 seconds, followed by 30 second pause, with active cooling via chilled air circulation to 4°C. SDS was added to 1% w/w, and the lysates were cleared by centrifugation (2 minutes, 1000 x G, 4°C), followed by spin filtration, with an initial pass through 1.2 µM filter (Acroprep Advance, No. 2022-05, PALL Corporation) followed by a second pass through 0.2 µM filter (Acroprep Advance, No. 2022-15, PALL Corporation), after which 6 µL of lysate was removed from each sample into a 96-well plate for the purposes of total protein determination by BCA assay. The remainder of each sample was frozen at -80°C.

*Protein Assay.* From the reserved 6 µL, each lysate was evaluated via BCA assay (Pierce) according to the manufacturer’s protocol. Triplicates at a 1:100 dilution and duplicates at a 1:200 dilution in PBS were incubated with 1X BCA reagent for one hour at 37°C. The samples were assayed for total protein concentration via absorbance at 562 nM in comparison to the standard curve comprised of serially diluted BSA.

*Peptide Preparation.* 50 µg of total protein (based on BCA assay) was removed from each sample to a new 96-well plate for subsequent processing. Sample volumes were normalized to 50 µL through the addition of the appropriate volume of lysis buffer. DTT was added to 5 mM final concentration in each sample, and samples were incubated at 37°C for 30 minutes. After equilibration to room temperature, iodoacetamide was added to 30 mM final concentration, and the samples were incubated at room temperature (RT) in the dark for 1 hour. Cysteine alkylation was quenched by addition of DTT to a final concentration of 25mM, and the samples were incubated in the dark for an additional 30 minutes at RT.

*Digestion.* Total protein was isolated via bead-based (SP3) precipitation protocol (3). Briefly, the magnetic beads were prepared by combining E3 and E7 Sera-Mag Carboxylate-Modified Magnetic SpeedBeads (Cytiva) at 1:1; vol:vol. Beads were washed three times with HPLC water prior to resuspension in lysis buffer for a final bead slurry concentration of 50 µg/µL. 15 µL of the pre-washed bead slurry and 100 µL 100% ethanol were added to each 50 µL sample containing 50 µg of protein. Proteins were allowed to bind and incubate with the bead slurry for 15 min at 25°C with gentle end-over-end rotation. Following protein binding, the supernatant was discarded, and the bead-captured proteins were washed three times with 80% ethanol and air dried. Purified protein-captured beads were resuspended in 100 mM EPPS, pH 8.0. Approximately 50 µg of protein/sample was digested at 25°C for 12 hours with lysyl endopeptidase (LysC, Wako Chemicals USA) at a 1:15; protease:protein (w/w) ratio. Following LysC digestion, the peptides were digested with trypsin at 37°C for 8 hours (Promega) at a 1:25; protease:protein (w/w) ratio. The digested peptides (supernatant) were purified and separated from the magnetic beads. Three post-digestion washes were performed to wash and ensure elution of all peptides from the beads with the following buffers: 0.2% formic acid, 80% acetonitrile/0.1% formic acid, and 95% acetonitrile/0.1% formic acid. With every wash and elution, the supernatant was transferred and combined with the initial supernatant and dried to completion by vacuum centrifugation. Digested samples were desalted via C18 microextraction (Waters, Product Code 186002318), and peptide eluates were dried via speedvac. Desalted peptides were resuspended in 10uL of 0.2% formic acid for LC-MS analysis.

*Preparation of potentially prion-infected for mass spectrometry.* For ki-3F4-FFI mice and matched controls, as a precaution in case spontaneously formed prions were present in the brain samples, lysis and sample processing were performed in the prion laboratory at McLaughlin Research Institute and only denatured peptide samples were sent for proteomics, consistent with best practices reported elsewhere (4, 5). Whole brain hemispheres were homogenized at 10% wt/vol in PBS (Gibco 14190) with 2% sodium dodecyl sulfate. A 25 µL aliquot, corresponding to 2.5 mg of tissue or an estimated ~250 µg of input protein, was used for peptide isolation. 25 µL PBS (Gibco 14190) was added, followed by TCEP (Thermo Fisher 77720) to 20 mM final, the solution was incubated at 37°C for 30 minutes, cooled to RT, then iodoacetamide (Sigma A3221) was added to 15 mM final, then incubated at room temperature 60 minutes. TCEP was added again to 40 mM final. Methanol/chloroform precipitation was performed with 100 µL chloroform, 400 µL methanol, and 300 µL water, and centrifugatoion at 3,214 x G for 10 minutes, followed by 2 washes in 200 µL methanol. The protein pellet was air-dried, 10 µL of guanidine hydrochloride was added, transferred to a new microcentrifuge tube, and incubated at 37°C for 30 minutes. 75 µL of 0.2M EPPS pH 8.1 (VWR AAJ612960AE) and 75 µL of HPLC water were added, followed by 5 µg of LysC (Promega V1671), then overnight incubation at room temperature, then 5 µg of trypsin (Promega V5113), and incubation at 37°C for 6 hours. Resulting peptides were frozen at -80°C before advancing to post-digestion wash and desalting steps described above.

*LC-MS Data Acquisition*. All data were acquired using an Orbitrap Eclipse mass spectrometer equipped with FAIMS-Pro, operating in nano-flow mode using a EASY nanoLC-1200 (ThermoFisher) system (Thermo). Separation was achieved using an in-house packed C18 Column (Sepax GP-C18 resin) that was 30 cm in length with 75 µm inner diameter. Peptides were separated over a gradient composed of Buffer B (80% acetonitrile/0.2% formic acid) in Buffer A (0.2% formic acid) was run according to the following table:

***Table S1. LC protocol used at IQ Proteomics.***

| **Time** | **duration (min)** | **flow (nL/min)** | **%B** |
| --- | --- | --- | --- |
| 0 | 0 | 300 | 10 |
| 50 | 50 | 300 | 40 |
| 55 | 5 | 300 | 100 |
| 60 | 5 | 300 | 100 |

Data was acquired via custom targeted assay with the instrument operating in tMS2 mode. Precursors were isolated using the quadrupole, and fragmented via HCD fragmentation. MS2 spectra were collected in the Orbitrap at 30K resolution. Maximum allowable injection time was set to 300ms.

***Table S2. Precursors in the IQ Proteomics assay.***

| **m/z** | **z** | **RT Time (min)** | **Window (min)** | **HCD Collision Energy (%)** | **FAIMS CV (V)** |
| --- | --- | --- | --- | --- | --- |
| 874.4391 | 2 | 27.5 | 4 | 26 | -40 |
| 877.4491 | 2 | 27.5 | 4 | 26 | -40 |
| 614.861 | 2 | 41 | 4 | 20 | -40 |
| 617.8711 | 2 | 41 | 4 | 20 | -40 |
| 444.2367 | 2 | 30 | 4 | 20 | -55 |
| 447.2579 | 2 | 30 | 4 | 20 | -55 |
| 454.2478 | 2 | 17 | 4 | 20 | -50 |
| 457.2579 | 2 | 17 | 4 | 20 | -50 |
| 721.3879 | 2 | 30 | 4 | 29 | -40 |
| 724.398 | 2 | 30 | 4 | 29 | -40 |
| 705.349 | 2 | 20 | 4 | 26 | -40 |
| 708.3591 | 2 | 20 | 4 | 26 | -40 |
| 674.367 | 2 | 30 | 4 | 23 | -40 |
| 677.377 | 2 | 30 | 4 | 23 | -40 |
| 444.2453 | 2 | 11 | 4 | 17 | -45 |
| 447.2553 | 2 | 11 | 4 | 17 | -45 |
| 961.4738 | 2 | 29 | 4 | 26 | -35 |
| 964.4838 | 2 | 29 | 4 | 26 | -35 |
| 629.8481 | 2 | 30 | 4 | 20 | -50 |
| 632.8582 | 2 | 30 | 4 | 20 | -50 |
| 883.4156 | 3 | 20 | 4 | 29 | -55 |
| 885.4223 | 3 | 20 | 4 | 29 | -55 |
| 502.2746 | 2 | 13 | 4 | 20 | -55 |
| 505.2846 | 2 | 13 | 4 | 20 | -55 |
| 756.8629 | 2 | 19 | 4 | 23 | -45 |
| 759.873 | 2 | 19 | 4 | 23 | -45 |
| 570.2644 | 2 | 14 | 4 | 23 | -50 |
| 573.2745 | 2 | 14 | 4 | 23 | -50 |
| 508.7722 | 2 | 5 | 8 | 23 | -65 |
| 511.7823 | 2 | 5 | 8 | 23 | -65 |
| 623.8193 | 2 | 39 | 4 | 26 | -50 |
| 626.8294 | 2 | 30 | 4 | 26 | -50 |
| 528.2902 | 2 | 29 | 4 | 20 | -50 |
| 531.3003 | 2 | 29 | 4 | 20 | -50 |
| 599.8423 | 2 | 25 | 4 | 20 | -40 |
| 596.8322 | 2 | 25 | 4 | 20 | -40 |
| 548.303 | 2 | 18 | 4 | 20 | -50 |
| 545.293 | 2 | 18 | 4 | 20 | -50 |
| 705.2925 | 2 | 31 | 6 | 23 | -45 |

*LC-MS Data Analysis*. LC-MS data was analyzed with the Skyline software package (www.skyline.ms). Peptide abundance was calculated as the ratio between the total area under the curve for the endogenous (light), chow-labeled (^13^C_6_ heavy) peptides. The following list of transitions was used:

***Table S3. Transitions used in the IQ Proteomics assay.***

| **Protein Name** | **Peptide Sequence** | **Precursor Mz** | **Precursor Charge** | **Product Mz** | **Product Charge** | **Fragment Ion** | **Isotope Label Type** |
| --- | --- | --- | --- | --- | --- | --- | --- |
| CAPZA2 | DIQDSLTVSNEVQTAK | 874.439063 | 2 | 1189.64229 | 1 | y11 | light |
| CAPZA2 | DIQDSLTVSNEVQTAK | 874.439063 | 2 | 1076.55823 | 1 | y10 | light |
| CAPZA2 | DIQDSLTVSNEVQTAK | 874.439063 | 2 | 975.510551 | 1 | y9 | light |
| CAPZA2 | DIQDSLTVSNEVQTAK | 874.439063 | 2 | 876.442137 | 1 | y8 | light |
| CAPZA2 | DIQDSLTVSNEVQTAK | 877.449127 | 2 | 1195.66242 | 1 | y11 | heavy |
| CAPZA2 | DIQDSLTVSNEVQTAK | 877.449127 | 2 | 1082.57836 | 1 | y10 | heavy |
| CAPZA2 | DIQDSLTVSNEVQTAK | 877.449127 | 2 | 981.53068 | 1 | y9 | heavy |
| CAPZA2 | DIQDSLTVSNEVQTAK | 877.449127 | 2 | 882.462266 | 1 | y8 | heavy |
| CRKL | TALALEVGDIVK | 614.861003 | 2 | 943.545873 | 1 | y9 | light |
| CRKL | TALALEVGDIVK | 614.861003 | 2 | 872.50876 | 1 | y8 | light |
| CRKL | TALALEVGDIVK | 614.861003 | 2 | 759.424696 | 1 | y7 | light |
| CRKL | TALALEVGDIVK | 614.861003 | 2 | 528.818607 | 2 | y10 | light |
| CRKL | TALALEVGDIVK | 614.861003 | 2 | 286.176132 | 1 | b3 | light |
| CRKL | TALALEVGDIVK | 617.871067 | 2 | 949.566002 | 1 | y9 | heavy |
| CRKL | TALALEVGDIVK | 617.871067 | 2 | 878.528889 | 1 | y8 | heavy |
| CRKL | TALALEVGDIVK | 617.871067 | 2 | 765.444825 | 1 | y7 | heavy |
| CRKL | TALALEVGDIVK | 617.871067 | 2 | 531.828671 | 2 | y10 | heavy |
| CRKL | TALALEVGDIVK | 617.871067 | 2 | 286.176132 | 1 | b3 | heavy |
| DC1L1 | LYGFPYK | 444.236721 | 2 | 774.382102 | 1 | y6 | light |
| DC1L1 | LYGFPYK | 444.236721 | 2 | 407.228896 | 1 | y3 | light |
| DC1L1 | LYGFPYK | 447.246786 | 2 | 780.402231 | 1 | y6 | heavy |
| DC1L1 | LYGFPYK | 447.246786 | 2 | 413.249025 | 1 | y3 | heavy |
| DCTN3 | ALLEGYNK | 454.247817 | 2 | 723.367181 | 1 | y6 | light |
| DCTN3 | ALLEGYNK | 454.247817 | 2 | 610.283117 | 1 | y5 | light |
| DCTN3 | ALLEGYNK | 454.247817 | 2 | 481.240524 | 1 | y4 | light |
| DCTN3 | ALLEGYNK | 457.257882 | 2 | 729.38731 | 1 | y6 | heavy |
| DCTN3 | ALLEGYNK | 457.257882 | 2 | 616.303246 | 1 | y5 | heavy |
| DCTN3 | ALLEGYNK | 457.257882 | 2 | 487.260653 | 1 | y4 | heavy |
| EMC7 | VLVDGEEHVGFLK | 721.387916 | 2 | 1229.61608 | 1 | y11 | light |
| EMC7 | VLVDGEEHVGFLK | 721.387916 | 2 | 1130.54766 | 1 | y10 | light |
| EMC7 | VLVDGEEHVGFLK | 721.387916 | 2 | 1015.52072 | 1 | y9 | light |
| EMC7 | VLVDGEEHVGFLK | 721.387916 | 2 | 615.311677 | 2 | y11 | light |
| EMC7 | VLVDGEEHVGFLK | 721.387916 | 2 | 565.77747 | 2 | y10 | light |
| EMC7 | VLVDGEEHVGFLK | 724.397981 | 2 | 1235.63621 | 1 | y11 | heavy |
| EMC7 | VLVDGEEHVGFLK | 724.397981 | 2 | 1136.56779 | 1 | y10 | heavy |
| EMC7 | VLVDGEEHVGFLK | 724.397981 | 2 | 1021.54085 | 1 | y9 | heavy |
| EMC7 | VLVDGEEHVGFLK | 724.397981 | 2 | 618.321742 | 2 | y11 | heavy |
| EMC7 | VLVDGEEHVGFLK | 724.397981 | 2 | 568.787535 | 2 | y10 | heavy |
| HMOX2 | ETLEDGLPVHDGK | 705.348988 | 2 | 1066.51636 | 1 | y10 | light |
| HMOX2 | ETLEDGLPVHDGK | 705.348988 | 2 | 937.473771 | 1 | y9 | light |
| HMOX2 | ETLEDGLPVHDGK | 705.348988 | 2 | 822.446828 | 1 | y8 | light |
| HMOX2 | ETLEDGLPVHDGK | 705.348988 | 2 | 652.3413 | 1 | y6 | light |
| HMOX2 | ETLEDGLPVHDGK | 705.348988 | 2 | 456.220122 | 1 | y4 | light |
| HMOX2 | ETLEDGLPVHDGK | 708.359052 | 2 | 1072.53649 | 1 | y10 | heavy |
| HMOX2 | ETLEDGLPVHDGK | 708.359052 | 2 | 943.4939 | 1 | y9 | heavy |
| HMOX2 | ETLEDGLPVHDGK | 708.359052 | 2 | 828.466957 | 1 | y8 | heavy |
| HMOX2 | ETLEDGLPVHDGK | 708.359052 | 2 | 658.361429 | 1 | y6 | heavy |
| HMOX2 | ETLEDGLPVHDGK | 708.359052 | 2 | 462.240251 | 1 | y4 | heavy |
| MAP2K1 | ISELGAGNGGVVFK | 674.366984 | 2 | 1018.56801 | 1 | y11 | light |
| MAP2K1 | ISELGAGNGGVVFK | 674.366984 | 2 | 905.483942 | 1 | y10 | light |
| MAP2K1 | ISELGAGNGGVVFK | 674.366984 | 2 | 848.462478 | 1 | y9 | light |
| MAP2K1 | ISELGAGNGGVVFK | 674.366984 | 2 | 777.425364 | 1 | y8 | light |
| MAP2K1 | ISELGAGNGGVVFK | 677.377048 | 2 | 1024.58814 | 1 | y11 | heavy |
| MAP2K1 | ISELGAGNGGVVFK | 677.377048 | 2 | 911.504071 | 1 | y10 | heavy |
| MAP2K1 | ISELGAGNGGVVFK | 677.377048 | 2 | 854.482607 | 1 | y9 | heavy |
| MAP2K1 | ISELGAGNGGVVFK | 677.377048 | 2 | 783.445493 | 1 | y8 | heavy |
| MAP7 | EQQILEK | 444.245275 | 2 | 630.382102 | 1 | y5 | light |
| MAP7 | EQQILEK | 444.245275 | 2 | 502.323525 | 1 | y4 | light |
| MAP7 | EQQILEK | 444.245275 | 2 | 389.239461 | 1 | y3 | light |
| MAP7 | EQQILEK | 444.245275 | 2 | 315.694689 | 2 | y5 | light |
| MAP7 | EQQILEK | 444.245275 | 2 | 386.167024 | 1 | b3 | light |
| MAP7 | EQQILEK | 447.255339 | 2 | 636.402231 | 1 | y5 | heavy |
| MAP7 | EQQILEK | 447.255339 | 2 | 508.343654 | 1 | y4 | heavy |
| MAP7 | EQQILEK | 447.255339 | 2 | 395.25959 | 1 | y3 | heavy |
| MAP7 | EQQILEK | 447.255339 | 2 | 318.704754 | 2 | y5 | heavy |
| MAP7 | EQQILEK | 447.255339 | 2 | 386.167024 | 1 | b3 | heavy |
| MARE3 | QGQDVAPPPNPGDQIFNK | 961.473771 | 2 | 1394.70629 | 1 | y13 | light |
| MARE3 | QGQDVAPPPNPGDQIFNK | 961.473771 | 2 | 1323.66918 | 1 | y12 | light |
| MARE3 | QGQDVAPPPNPGDQIFNK | 961.473771 | 2 | 1226.61641 | 1 | y11 | light |
| MARE3 | QGQDVAPPPNPGDQIFNK | 961.473771 | 2 | 1129.56365 | 1 | y10 | light |
| MARE3 | QGQDVAPPPNPGDQIFNK | 961.473771 | 2 | 918.467957 | 1 | y8 | light |
| MARE3 | QGQDVAPPPNPGDQIFNK | 964.483836 | 2 | 1400.72642 | 1 | y13 | heavy |
| MARE3 | QGQDVAPPPNPGDQIFNK | 964.483836 | 2 | 1329.68931 | 1 | y12 | heavy |
| MARE3 | QGQDVAPPPNPGDQIFNK | 964.483836 | 2 | 1232.63654 | 1 | y11 | heavy |
| MARE3 | QGQDVAPPPNPGDQIFNK | 964.483836 | 2 | 1135.58378 | 1 | y10 | heavy |
| MARE3 | QGQDVAPPPNPGDQIFNK | 964.483836 | 2 | 924.488086 | 1 | y8 | heavy |
| NCAM2 | EVLNAETIEIK | 629.848092 | 2 | 1030.5779 | 1 | y9 | light |
| NCAM2 | EVLNAETIEIK | 629.848092 | 2 | 917.493838 | 1 | y8 | light |
| NCAM2 | EVLNAETIEIK | 629.848092 | 2 | 803.45091 | 1 | y7 | light |
| NCAM2 | EVLNAETIEIK | 629.848092 | 2 | 732.413797 | 1 | y6 | light |
| NCAM2 | EVLNAETIEIK | 632.858157 | 2 | 1036.59803 | 1 | y9 | heavy |
| NCAM2 | EVLNAETIEIK | 632.858157 | 2 | 923.513967 | 1 | y8 | heavy |
| NCAM2 | EVLNAETIEIK | 632.858157 | 2 | 809.471039 | 1 | y7 | heavy |
| NCAM2 | EVLNAETIEIK | 632.858157 | 2 | 738.433926 | 1 | y6 | heavy |
| NCAM2 | VSNDIIQSK | 502.274563 | 2 | 904.473437 | 1 | y8 | light |
| NCAM2 | VSNDIIQSK | 502.274563 | 2 | 817.441408 | 1 | y7 | light |
| NCAM2 | VSNDIIQSK | 502.274563 | 2 | 588.371538 | 1 | y5 | light |
| NCAM2 | VSNDIIQSK | 502.274563 | 2 | 475.287474 | 1 | y4 | light |
| NCAM2 | VSNDIIQSK | 505.284628 | 2 | 910.493566 | 1 | y8 | heavy |
| NCAM2 | VSNDIIQSK | 505.284628 | 2 | 823.461537 | 1 | y7 | heavy |
| NCAM2 | VSNDIIQSK | 505.284628 | 2 | 594.391667 | 1 | y5 | heavy |
| NCAM2 | VSNDIIQSK | 505.284628 | 2 | 481.307603 | 1 | y4 | heavy |
| PRNP | VVEQMC[+57]VTQYQK | 756.862894 | 2 | 1057.48051 | 1 | y8 | light |
| PRNP | VVEQMC[+57]VTQYQK | 756.862894 | 2 | 926.440028 | 1 | y7 | light |
| PRNP | VVEQMC[+57]VTQYQK | 756.862894 | 2 | 667.340966 | 1 | y5 | light |
| PRNP | VVEQMC[+57]VTQYQK | 759.872958 | 2 | 1063.50064 | 1 | y8 | heavy |
| PRNP | VVEQMC[+57]VTQYQK | 759.872958 | 2 | 932.460157 | 1 | y7 | heavy |
| PRNP | VVEQMC[+57]VTQYQK | 759.872958 | 2 | 673.361095 | 1 | y5 | heavy |
| PRNP | GENFTETDVK | 570.264393 | 2 | 953.457452 | 1 | y8 | light |
| PRNP | GENFTETDVK | 570.264393 | 2 | 839.414525 | 1 | y7 | light |
| PRNP | GENFTETDVK | 570.264393 | 2 | 692.346111 | 1 | y6 | light |
| PRNP | GENFTETDVK | 570.264393 | 2 | 591.298432 | 1 | y5 | light |
| PRNP | GENFTETDVK | 573.274457 | 2 | 959.477581 | 1 | y8 | heavy |
| PRNP | GENFTETDVK | 573.274457 | 2 | 845.434654 | 1 | y7 | heavy |
| PRNP | GENFTETDVK | 573.274457 | 2 | 698.36624 | 1 | y6 | heavy |
| PRNP | GENFTETDVK | 573.274457 | 2 | 597.318561 | 1 | y5 | heavy |
| PRNP | QHTVTTTTK | 508.772188 | 2 | 751.41961 | 1 | y7 | light |
| PRNP | QHTVTTTTK | 508.772188 | 2 | 650.371932 | 1 | y6 | light |
| PRNP | QHTVTTTTK | 508.772188 | 2 | 450.255839 | 1 | y4 | light |
| PRNP | QHTVTTTTK | 508.772188 | 2 | 349.208161 | 1 | y3 | light |
| PRNP | QHTVTTTTK | 511.782252 | 2 | 757.439739 | 1 | y7 | heavy |
| PRNP | QHTVTTTTK | 511.782252 | 2 | 656.392061 | 1 | y6 | heavy |
| PRNP | QHTVTTTTK | 511.782252 | 2 | 456.275968 | 1 | y4 | heavy |
| PRNP | QHTVTTTTK | 511.782252 | 2 | 355.22829 | 1 | y3 | heavy |
| SCRN1 | DEAWVLETVGK | 623.819335 | 2 | 1002.56186 | 1 | y9 | light |
| SCRN1 | DEAWVLETVGK | 623.819335 | 2 | 931.524744 | 1 | y8 | light |
| SCRN1 | DEAWVLETVGK | 623.819335 | 2 | 745.445431 | 1 | y7 | light |
| SCRN1 | DEAWVLETVGK | 623.819335 | 2 | 646.377017 | 1 | y6 | light |
| SCRN1 | DEAWVLETVGK | 623.819335 | 2 | 502.193239 | 1 | b4 | light |
| SCRN1 | DEAWVLETVGK | 626.8294 | 2 | 1008.58199 | 1 | y9 | heavy |
| SCRN1 | DEAWVLETVGK | 626.8294 | 2 | 937.544873 | 1 | y8 | heavy |
| SCRN1 | DEAWVLETVGK | 626.8294 | 2 | 751.46556 | 1 | y7 | heavy |
| SCRN1 | DEAWVLETVGK | 626.8294 | 2 | 652.397146 | 1 | y6 | heavy |
| SCRN1 | DEAWVLETVGK | 626.8294 | 2 | 502.193239 | 1 | b4 | heavy |
| UBC12 | DINELNLPK | 528.290213 | 2 | 827.462144 | 1 | y7 | light |
| UBC12 | DINELNLPK | 528.290213 | 2 | 713.419216 | 1 | y6 | light |
| UBC12 | DINELNLPK | 528.290213 | 2 | 584.376623 | 1 | y5 | light |
| UBC12 | DINELNLPK | 528.290213 | 2 | 471.292559 | 1 | y4 | light |
| UBC12 | DINELNLPK | 528.290213 | 2 | 414.23471 | 2 | y7 | light |
| UBC12 | DINELNLPK | 531.300278 | 2 | 833.482273 | 1 | y7 | heavy |
| UBC12 | DINELNLPK | 531.300278 | 2 | 719.439345 | 1 | y6 | heavy |
| UBC12 | DINELNLPK | 531.300278 | 2 | 590.396752 | 1 | y5 | heavy |
| UBC12 | DINELNLPK | 531.300278 | 2 | 477.312688 | 1 | y4 | heavy |
| UBC12 | DINELNLPK | 531.300278 | 2 | 417.244774 | 2 | y7 | heavy |
| VIGLN | LGQALTEVYAK | 596.832245 | 2 | 1079.57315 | 1 | y10 | light |
| VIGLN | LGQALTEVYAK | 596.832245 | 2 | 894.49311 | 1 | y8 | light |
| VIGLN | LGQALTEVYAK | 596.832245 | 2 | 609.324253 | 1 | y5 | light |
| VIGLN | LGQALTEVYAK | 596.832245 | 2 | 480.28166 | 1 | y4 | light |
| VIGLN | LGQALTEVYAK | 596.832245 | 2 | 511.779482 | 2 | y9 | light |
| VIGLN | LGQALTEVYAK | 599.84231 | 2 | 1085.59328 | 1 | y10 | heavy |
| VIGLN | LGQALTEVYAK | 599.84231 | 2 | 900.513239 | 1 | y8 | heavy |
| VIGLN | LGQALTEVYAK | 599.84231 | 2 | 615.344382 | 1 | y5 | heavy |
| VIGLN | LGQALTEVYAK | 599.84231 | 2 | 486.301789 | 1 | y4 | heavy |
| VIGLN | LGQALTEVYAK | 599.84231 | 2 | 514.789546 | 2 | y9 | heavy |
| WDR47 | VTDLQGDLTK | 545.292953 | 2 | 990.510216 | 1 | y9 | light |
| WDR47 | VTDLQGDLTK | 545.292953 | 2 | 661.351531 | 1 | y6 | light |
| WDR47 | VTDLQGDLTK | 545.292953 | 2 | 316.150312 | 1 | b3 | light |
| WDR47 | VTDLQGDLTK | 548.303018 | 2 | 996.530345 | 1 | y9 | heavy |
| WDR47 | VTDLQGDLTK | 548.303018 | 2 | 667.37166 | 1 | y6 | heavy |
| WDR47 | VTDLQGDLTK | 548.303018 | 2 | 316.150312 | 1 | b3 | heavy |

*Lower limit of quantification (LLQ) sample preparation.* Internal standard peptides labeled with ^13^C_6_ ^15^N_2_ lysine (Biosynth, Gardner MA, USA) were mixed into an equimolar solution, and a 12-point, two-fold serial dilution of the peptide mix was prepared by diluting into 0.2% formic acid solution such that the following amounts of peptide were evaluated by LC-MS according to Table S4.

***Table S4. Synthetic peptide serial dilutions to determine lower limit of quantification (LLQ)***

| **curve point** | **fmol peptide on column** |
| --- | --- |
| 12 | 40 |
| 11 | 20 |
| 10 | 10 |
| 9 | 5 |
| 8 | 2.5 |
| 7 | 1.25 |
| 6 | 0.625 |
| 5 | 0.313 |
| 4 | 0.156 |
| 3 | 0.0781 |
| 2 | 0.039 |
| 1 | 0.0195 |

*LLQ LC-MS Runs.* LC and MS instrument parameters for LoQ runs were identical to those for the study samples with the exception that masses corresponding to the heavy synthetic peptide standards (containing terminal ^13^C_6_ ^15^N_2_ lysine) were evaluated:

***Table S4. Precursors for LLQ determination.***

| m/z | z | RT Time (min) | Window (min) | HCD Collision Energy (%) | FAIMS CV (V) |
| --- | --- | --- | --- | --- | --- |
| 874.4391 | 2 | 27.7 | 10 | 26 | -40 |
| 878.4462 | 2 | 27.7 | 10 | 26 | -40 |
| 614.861 | 2 | 40.2 | 10 | 20 | -40 |
| 618.8681 | 2 | 40.2 | 10 | 20 | -40 |
| 444.2367 | 2 | 31.6 | 10 | 20 | -55 |
| 448.2438 | 2 | 31.6 | 10 | 20 | -55 |
| 454.2478 | 2 | 20.3 | 10 | 20 | -50 |
| 458.2549 | 2 | 20.3 | 10 | 20 | -50 |
| 721.3879 | 2 | 30.9 | 10 | 29 | -40 |
| 725.395 | 2 | 30.9 | 10 | 29 | -40 |
| 705.349 | 2 | 22.1 | 10 | 26 | -40 |
| 709.3561 | 2 | 22.1 | 10 | 26 | -40 |
| 674.367 | 2 | 30.5 | 10 | 23 | -40 |
| 678.3741 | 2 | 30.5 | 10 | 23 | -40 |
| 444.2453 | 2 | 13.4 | 10 | 17 | -45 |
| 448.2524 | 2 | 13.4 | 10 | 17 | -45 |
| 961.4738 | 2 | 29.5 | 10 | 26 | -35 |
| 965.4809 | 2 | 29.5 | 10 | 26 | -35 |
| 629.8481 | 2 | 31 | 10 | 20 | -50 |
| 633.8552 | 2 | 31 | 10 | 20 | -50 |
| 883.4156 | 3 | 21.6 | 10 | 29 | -55 |
| 886.087 | 3 | 21.6 | 10 | 29 | -55 |
| 502.2746 | 2 | 15.2 | 10 | 20 | -55 |
| 506.2817 | 2 | 15.2 | 10 | 20 | -55 |
| 400.2554 | 2 | 30.7 | 10 | 20 | -60 |
| 405.2596 | 2 | 30.7 | 10 | 20 | -60 |
| 522.7409 | 2 | 10 | 10 | 20 | -55 |
| 527.745 | 2 | 10 | 10 | 20 | -55 |
| 778.3658 | 2 | 27.4 | 10 | 23 | -40 |
| 783.3699 | 2 | 27.4 | 10 | 23 | -40 |
| 544.7358 | 2 | 10.8 | 10 | 20 | -60 |
| 549.7399 | 2 | 10.8 | 10 | 20 | -60 |
| 756.8629 | 2 | 21.1 | 10 | 23 | -45 |
| 760.87 | 2 | 21.1 | 10 | 23 | -45 |
| 570.2644 | 2 | 16 | 10 | 23 | -50 |
| 574.2715 | 2 | 16 | 10 | 23 | -50 |
| 508.7722 | 2 | 7.5 | 10 | 23 | -65 |
| 512.7793 | 2 | 7.5 | 10 | 23 | -65 |
| 545.2572 | 2 | 8.8 | 10 | 32 | -60 |
| 550.2614 | 2 | 8.8 | 10 | 32 | -60 |
| 623.8193 | 2 | 37.9 | 10 | 26 | -50 |
| 627.8264 | 2 | 37.9 | 10 | 26 | -50 |
| 483.238 | 2 | 18 | 10 | 26 | -50 |
| 488.2421 | 2 | 18 | 10 | 26 | -50 |
| 525.8084 | 2 | 43.5 | 10 | 20 | -55 |
| 530.8125 | 2 | 43.5 | 10 | 20 | -55 |
| 528.2902 | 2 | 29.7 | 10 | 20 | -50 |
| 532.2973 | 2 | 29.7 | 10 | 20 | -50 |
| 596.8322 | 2 | 26.7 | 10 | 20 | -40 |
| 600.8393 | 2 | 26.7 | 10 | 20 | -40 |
| 545.293 | 2 | 19.7 | 10 | 20 | -50 |
| 549.3001 | 2 | 19.7 | 10 | 20 | -50 |

***Table S6. Fragment ions corresponding to the synthetic peptides for LLQ determination.***

| Peptide | Precursor m/z | Fragment m/z | Fragment Ion |
| --- | --- | --- | --- |
| DIQDSLTVSNEVQTAK | 878.446162 | 1197.65649 | y11 |
| DIQDSLTVSNEVQTAK | 878.446162 | 1084.57243 | y10 |
| DIQDSLTVSNEVQTAK | 878.446162 | 983.52475 | y9 |
| DIQDSLTVSNEVQTAK | 878.446162 | 884.456336 | y8 |
| TALALEVGDIVK | 618.868102 | 951.560072 | y9 |
| TALALEVGDIVK | 618.868102 | 880.522959 | y8 |
| TALALEVGDIVK | 618.868102 | 767.438895 | y7 |
| TALALEVGDIVK | 618.868102 | 532.825706 | y10 |
| LYGFPYK | 448.243821 | 782.396301 | y6 |
| LYGFPYK | 448.243821 | 619.332973 | y5 |
| LYGFPYK | 448.243821 | 562.311509 | y4 |
| LYGFPYK | 448.243821 | 415.243095 | y3 |
| LYGFPYK | 448.243821 | 391.701789 | y6 |
| ALLEGYNK | 458.254917 | 731.38138 | y6 |
| ALLEGYNK | 458.254917 | 618.297316 | y5 |
| ALLEGYNK | 458.254917 | 489.254723 | y4 |
| VLVDGEEHVGFLK | 725.395016 | 1237.63028 | y11 |
| VLVDGEEHVGFLK | 725.395016 | 1138.56186 | y10 |
| VLVDGEEHVGFLK | 725.395016 | 1023.53492 | y9 |
| VLVDGEEHVGFLK | 725.395016 | 619.318777 | y11 |
| VLVDGEEHVGFLK | 725.395016 | 569.78457 | y10 |
| ETLEDGLPVHDGK | 709.356087 | 1074.53056 | y10 |
| ETLEDGLPVHDGK | 709.356087 | 945.48797 | y9 |
| ETLEDGLPVHDGK | 709.356087 | 830.461027 | y8 |
| ETLEDGLPVHDGK | 709.356087 | 660.355499 | y6 |
| ETLEDGLPVHDGK | 709.356087 | 464.234321 | y4 |
| ISELGAGNGGVVFK | 678.374083 | 1026.58221 | y11 |
| ISELGAGNGGVVFK | 678.374083 | 913.498141 | y10 |
| ISELGAGNGGVVFK | 678.374083 | 856.476677 | y9 |
| ISELGAGNGGVVFK | 678.374083 | 785.439563 | y8 |
| EQQILEK | 448.252374 | 638.396301 | y5 |
| EQQILEK | 448.252374 | 510.337724 | y4 |
| EQQILEK | 448.252374 | 397.25366 | y3 |
| EQQILEK | 448.252374 | 319.701789 | y5 |
| EQQILEK | 448.252374 | 386.167024 | b3 |
| QGQDVAPPPNPGDQIFNK | 965.480871 | 1402.72049 | y13 |
| QGQDVAPPPNPGDQIFNK | 965.480871 | 1331.68338 | y12 |
| QGQDVAPPPNPGDQIFNK | 965.480871 | 1234.63061 | y11 |
| QGQDVAPPPNPGDQIFNK | 965.480871 | 1137.57785 | y10 |
| QGQDVAPPPNPGDQIFNK | 965.480871 | 926.482156 | y8 |
| EVLNAETIEIK | 633.855192 | 1038.5921 | y9 |
| EVLNAETIEIK | 633.855192 | 925.508037 | y8 |
| EVLNAETIEIK | 633.855192 | 811.465109 | y7 |
| EVLNAETIEIK | 633.855192 | 740.427996 | y6 |
| EVLNAETIEIK | 633.855192 | 611.385402 | y5 |
| ITNHEDGSPVNEPNETTPLTEPEK | 886.086987 | 1363.6831 | y12 |
| ITNHEDGSPVNEPNETTPLTEPEK | 886.086987 | 821.449459 | y7 |
| ITNHEDGSPVNEPNETTPLTEPEK | 886.086987 | 1293.57059 | b12 |
| VSNDIIQSK | 506.281663 | 912.487636 | y8 |
| VSNDIIQSK | 506.281663 | 825.455607 | y7 |
| VSNDIIQSK | 506.281663 | 711.41268 | y6 |
| VSNDIIQSK | 506.281663 | 596.385737 | y5 |
| VSNDIIQSK | 506.281663 | 483.301673 | y4 |
| VVEQMC[+57.021464]VTQYQK | 760.869993 | 1065.49471 | y8 |
| VVEQMC[+57.021464]VTQYQK | 760.869993 | 934.454227 | y7 |
| VVEQMC[+57.021464]VTQYQK | 760.869993 | 675.355165 | y5 |
| GENFTETDVK | 574.271492 | 961.471651 | y8 |
| GENFTETDVK | 574.271492 | 847.428724 | y7 |
| GENFTETDVK | 574.271492 | 700.36031 | y6 |
| GENFTETDVK | 574.271492 | 599.312631 | y5 |
| QHTVTTTTK | 512.779287 | 759.433809 | y7 |
| QHTVTTTTK | 512.779287 | 658.386131 | y6 |
| QHTVTTTTK | 512.779287 | 559.317717 | y5 |
| QHTVTTTTK | 512.779287 | 458.270038 | y4 |
| QHTVTTTTK | 512.779287 | 357.22236 | y3 |
| DEAWVLETVGK | 627.826435 | 1010.57606 | y9 |
| DEAWVLETVGK | 627.826435 | 939.538943 | y8 |
| DEAWVLETVGK | 627.826435 | 753.45963 | y7 |
| DEAWVLETVGK | 627.826435 | 654.391216 | y6 |
| DEAWVLETVGK | 627.826435 | 502.193239 | b4 |
| DINELNLPK | 532.297313 | 835.476343 | y7 |
| DINELNLPK | 532.297313 | 721.433415 | y6 |
| DINELNLPK | 532.297313 | 592.390822 | y5 |
| DINELNLPK | 532.297313 | 479.306758 | y4 |
| DINELNLPK | 532.297313 | 418.241809 | y7 |
| LGQALTEVYAK | 600.839345 | 1087.58735 | y10 |
| LGQALTEVYAK | 600.839345 | 902.507309 | y8 |
| LGQALTEVYAK | 600.839345 | 831.470195 | y7 |
| LGQALTEVYAK | 600.839345 | 718.386131 | y6 |
| LGQALTEVYAK | 600.839345 | 515.786581 | y9 |
| VTDLQGDLTK | 549.300053 | 998.524415 | y9 |
| VTDLQGDLTK | 549.300053 | 897.476737 | y8 |
| VTDLQGDLTK | 549.300053 | 669.36573 | y6 |
| VTDLQGDLTK | 549.300053 | 541.307152 | y5 |
| VTDLQGDLTK | 549.300053 | 316.150312 | b3 |

***Targeted mass spectrometry at Charles River Labs.***

*Tissue homogenization.* Colon and brain tissue were homogenized in TPER reagent (Thermo No. 78510) with 1X HALT protease inhibitor (Thermo No. 78430) at a ratio of 1 g tissue to 9 mL buffer.

*Total protein estimation by BCA assay.* 10 µL aliquots of tissue homogenates (see above) were diluted 10-fold in water for total protein estimation. Seven calibration standards concentrations (2, 1, 0.5, 0.25, 0.125, 0.0625, 0.03125) were prepared with serial 2x dilution in milliQ water. BCA reagent was prepared by mixing reagent A and B in a 50:1 ratio. 20 µL of sample and 200 µL of BCA reagent were added to each well of a clear bottom 96 well plate and incubated at 37°C for 20 minutes. Absorbance of each well was read at 562 nm using a UV spectrometer. Samples were diluted to 3 mg/mL total protein concentration based on BCA assay data.

*Extraction procedure.* 60 µL of mixed reagent consisting of 20 µL bovine serum albumin (BSA), 30 µL ammonium bicarbonate (ABC, 100 mM), and 10 µL dithiothreitol (DTT, 250 mM) was added to 80 µL of the 3 mg/mL sample and mixed before incubating at 95°C for 10 minutes at 300 rpm shaking. Samples were then allowed to cool for 10 minutes at room temperature. 10 µL of iodoacetamide (IAA, 500 mM) was added to the samples, vortexed to mix and incubated at RT in the dark for 30 minutes. 1 mL of cold acetone was added to the samples and vortexed to mix again prior to incubating at -80°C for 30 minutes. Samples were spun at 16,000 x G for 5 minutes and supernatant was discarded. The protein pellet was dried in the hood, washed with 500 µL cold methanol and dried for 30 additional minutes in the hood at room temperature. The protein pellet was resuspended in 60 µL of 50 mM ABC plus 20 µL of diluted trypsin (0.16 µg/µL) for a final quantity of 3.2 µg trypsin to 240 µg protein (1:75 enzyme:protein ratio) and mixed with a pipette before incubating for 16 hours at 37°C. An internal standard (IS) of peptide with isotopically labeled lysine (^13^C, ^15^N) custom synthesized by Vivitide was used. The reaction was stopped by adding 10 µL of IS (prepared in 0.5% FA ACN-H_2_O 80:20) and 10 µL FA in ACN:H_2_O (80:20) and vortexing to mix before centrifugation at 8,000 x G for 5 minutes. Supernatants were transferred to LCMS vials.

*Calibration curve.* 80 µL of 50 mM ammonium bicarbonate was aliquoted into the tubes designated for blanks and standards. The calibration curve was prepared with light peptides standard custom synthesized by Vivitide. 10 µL of ACN:H2O:FA 1:10:1 was aliquoted into each 80 µL sample, blank, or standard. 10 µL of standards were added to tubes labeled for standards. 10 µL of IS in 80:20:0.5 ACN:H_2_O:FA was added to all standard, QC, and blank tubes except for double blank tubes. 10 µL of 80:20:0.5 ACN:H_2_O:FA was added to double blanks. All Eppendorf tubes were covered and vortexed prior to being centrifuged for 5 minutes at 8000 x G. At least 80 µL of the samples were transferred into a clean 96-well plate.
